# Statistical analysis of spatially resolved transcriptomic data by incorporating multi-omics auxiliary information

**DOI:** 10.1101/2022.04.22.489194

**Authors:** Yan Li, Xiang Zhou, Hongyuan Cao

## Abstract

Effective control of false discovery rate is key for multiplicity problems. Here, we consider incorporating informative covariates from external datasets in the multiple testing procedure to boost statistical power while maintaining false discovery rate control. In particular, we focus on the statistical analysis of innovative high-dimensional spatial transcriptomic data while incorporating external multi-omics data that provide distinct but complementary information to the detection of spatial expression patterns. We extend OrderShapeEM, an efficient covariate-assisted multiple testing procedure that incorporates one auxiliary study, to make it permissible to incorporate multiple external omics studies, to boost statistical power of spatial expression pattern detection. Specifically, we first use a recently proposed computationally efficient statistical analysis method, spatial pattern recognition via kernels, to produce the primary test statistics for spatial transcriptomic data. Afterwards, we construct the auxiliary covariate by combining information from multiple external omics studies, such as bulk or single-cell RNA-seq data and genome wise association study data, using the Cauchy combination rule. Finally, we extend and implement the integrative analysis method OrderShapeEM on the primary *p*-values along with auxiliary data incorporating multi-omics information for efficient covariate-assisted spatial expression analysis. We conduct a series of realistic simulations to evaluate the performance of our method with known ground truth. Four case studies in mouse olfactory bulb, mouse cerebellum, human breast cancer and human heart tissues further demonstrate the substantial power gain of our method in detecting genes with spatial expression patterns compared to existing classic approaches that do not utilize any external information.

## 1 Introduction

Modern advancement of science and technology has enabled gene-expression profiling with spatial location information on tissues or cell cultures (Stahl *et al*., 2016; Stickels *et al*., 2021). Such data allow scientists to study the spatial organization of the transcriptome on tissue sections or across cells on a cell culture, providing substantial insights into the complex biological phenomena. As a first step to characterizing the transcriptomic landscape of tissues, identifying genes that display spatial expression (SE) patterns is vitally important and can provide crucial and meaningful insights in the fields of embryology, oncology, immunology and histology (Asp *et al*., 2020). Because SE analysis tests tens of thousands of genes simultaneously, an acute problem is multiple comparison adjustment. A classic approach for multiple comparison is the false discovery rate (FDR) control, where it is assumed that each gene has equal likelihood to display SE pattern *a priori* (Benjamini and Hochberg, 1995). However, other omics datasets from external studies (i.e. genomics, epigenomics and proteomics, etc.) often collect additional genomics information in the same or similar tissue and can provide distinct but complementary information to statistical analysis of spatially resolved transcriptomic data. Leveraging such mutually informative but partially independent auxiliary information from external datasets will likely lead to discovery of more, novel, functional and reproducible features.

Statistical analysis of spatially resolved transcriptomic data requires proper modeling of the spatial correlation structure in the gene expression measurements collected across spatial locations. For SE analysis, Trendsceek (Edsgard *et al*., 2018) and SpatialDE (Svensson *et al*., 2018) transform raw count data into normalized expression data, and develop statistical tests after the normalization. Spatial pattern recognition via kernels (SPARK) (Sun *et al*., 2020) directly models the raw read counts with a generalized linear spatial model (GLSM) and produces well-calibrated *p*-values via a score test. Under the null hypothesis that a particular gene does not exhibit any SE pattern, the GLSM is reduced to the generalized linear mixed model, and the parameters estimated from the null model are used to construct score test statistics. Once *p*-values are obtained, a multiple comparison procedure that allows for arbitrary dependence across genes such as BY procedure (Benjamini and Yekutieli, 2001) is used to conclude statistical significance.

As a multiple comparison criterion, the FDR is defined as the expectation of the number of false rejections over total number of rejections (Benjamini and Hochberg, 1995). At a pre-specified FDR level *α*, the actual control level of the BH step-up procedure is π_0_*α*, where π_0_ is the proportion of true null hypotheses (Benjamini and Hochberg, 1995). The *q*-value procedure (Storey and Tibshirani, 2003) plugs in a conservative estimate of π_0_ to improve the power of BH. The BH and *q*-value procedures assume that hypotheses are exchangeable. Recently, a plethora of methods have been developed to incorporate side information into FDR control. Basically, with auxiliary information, genes have different probability to be SE *a priori*. For genes that are more likely to be SE *a priori*, the *p*-value threshold could be loosened; on the other hand, for genes that are less likely to be SE *a priori*, the *p*-value threshold is tightened. Such flexible modeling controls the overall FDR and improves statistical power. For instance, independent hypothesis weighting (IHW) divides hypotheses into groups based on the covariate and all hypotheses within a group have the same weight. Then a weighted BH procedure is used to achieve an overall FDR control (Genovese *et al*., 2006; Ignatiadis *et al*., 2016; Ignatiadis and Huber, 2021). SABHA (Li and Barber, 2019) incorporates prior information about the location pattern of signals and nulls, and uses a data-adaptive weight to reweigh the *p*-values. AdaPT (Lei and Fithian, 2018) iteratively estimates the *p*-value threshold using partially masked *p*-values. The false discovery proportion is estimated by virtue of the symmetric distribution of *p*-values under the null hypothesis. Most recently, Cao *et al*. (2022) proposes a general and efficient framework OrderShapeEM to incorporate auxiliary information of one dimension. OrderShapeEM uses the local false discovery rate (Lfdr) as the test statistic and adaptively estimates the prior probabilities of being null in Lfdr incorporating auxiliary information (Sun and Cai, 2007; Efron, 2008; Cao *et al*., 2013). It imposes a monotone constraint on the prior probabilities of being null and the *p*-values, which is tuning-parameter free and robust to mis-specification of prior information. In terms of implementation, it utilizes the EM algorithm and pool-adjacent-violator-algorithm (PAVA) for fast computation (Dempster *et al*., 1977; Robertson *et al*., 1988).

In SE analysis, in addition to the primary spatial transcriptomic data, there is a wealth of omics data that can provide useful auxiliary information. For example, integrating spatial transcriptomic data with scRNA-seq data may increase our understanding of the role of specific cell sub-populations and their interactions in development, homeostasis and disease (Longo *et al*., 2021). How to effectively exploit such auxiliary information to improve the statistical power of SE analysis has not been well explored. In this paper, by extending OrderShapeEM, we aim to conduct an integrative analysis with spatially resolved transcriptomic data using SPARK incorporating multi-omics auxiliary information.

Computationally, OrderShapeEM does not involve any tuning parameter and the implementation is scalable with built-in PAVA and EM algorithms. Statistically speaking, signals that appear not only in spatially resolved transcriptomic data but also in other omics data are more likely to be true signals and the proposed method can reduce false positive findings as fewer of them may arise by chance in multiple conditions. On the other hand, many SE genes have only moderate-to-weak effects in a single study, and would be missed using standard spatial transcriptomic data analysis after stringent multiple comparison. The integrative analysis framework borrows rich and complementary information across different studies, which can substantially improve statistical power. For auxiliary information, we only need summary level statistics, such as *p*-values, which can preserve the confidentiality and individual privacy. We impose a monotone relationship between the primary *p*-values from spatial transcriptomic data and the auxiliary covariate sequence. The scientific rationale behind this monotone constraint is pleiotropy which indicates that one gene can be associated with many different traits (Solovieff *et al*., 2013). For a gene exhibiting pleiotropy in the primary study and the auxiliary study, the *p*-values are more likely to be small in both studies, rendering the monotone relationship between them. Importantly, OrderShapeEM is robust to mis-specification of the order structure provided by auxiliary information. This flexibility makes our method have comparable power with conventional method that does not incorporate covariate information when the covariate is uninformative or weakly informative. One of the limitations of the original OrderShapeEM is that only scalar covariate can be incorporated. We broaden the application scope to allow multiple omics data as auxiliary information. This is achieved through a recently developed Cauchy combination rule that combines multiple *p*-values under arbitrary dependence into one *p*-value (Liu *et al*., 2019; Liu and Xie, 2020).

As a proof-of-concept, we implemented the proposed method to four spatially resolved transcriptomics datasets and identified additional SE genes that were missed by the original SPARK method. We found that incorporating multiple auxiliary datasets leads to more discoveries than incorporating one auxiliary dataset. Many of the additionally identified SE genes can be validated through several reference gene lists and have plausible biological implications.

## 2 Materials and methods

We extend an integrative analysis method, OrderShapeEM (Cao *et al*., 2022), to detect SE genes from spatially resolved transcriptomic data incorporating multi-omics auxiliary data. We remark that OrderShapeEM is a generic multiple comparison procedure with auxiliary information that is applicable to many types of omics data. OrderShapeEM takes as input *m p*-values 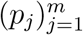 from the primary study and a corresponding covariate sequence 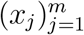 from auxiliary information, and calculates the Lfdr based on an empirical Bayesian two-group mixture model (Efron, 2008). Unknown parameters and functions in the two-group mixture model are estimated by the combination of PAVA and EM algorithms. A step-up procedure for Lfdr is used to calculate the threshold for rejection and outputs a binary indicator signifying whether certain gene is SE or not. See Fig. 1 for the full pipeline of our approach. In the following, we first describe the modeling of spatial transcriptomic data, then propose a way to combine auxiliary information from multiple studies, and finally incorporate the combination of multi-omics data into the primary multiple testing procedure with OrderShapeEM.

**Figure 1:**
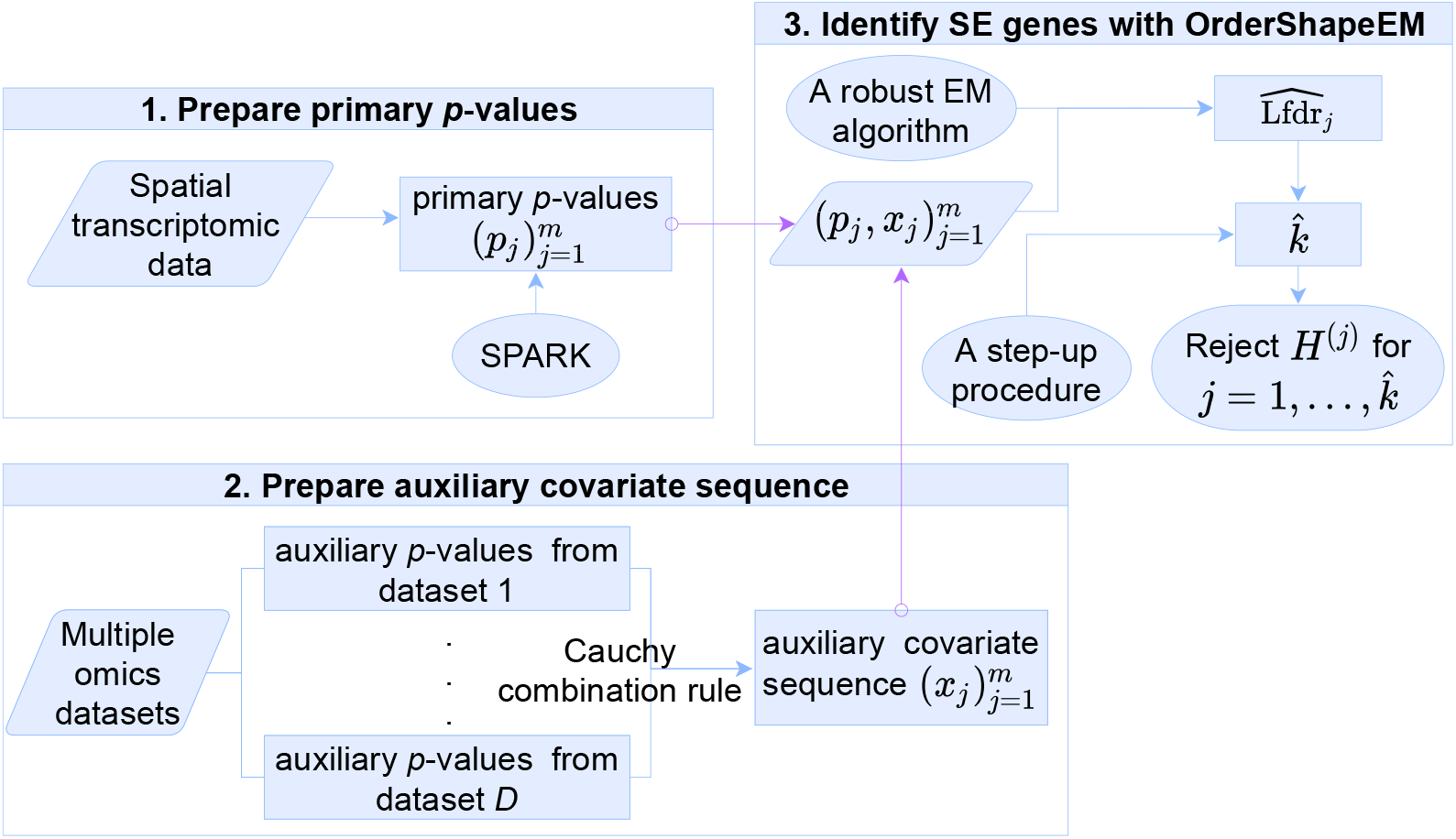
Schematic plot of the proposed integrative analysis.

### 2.1 Primary *p*-value construction with SPARK

Our main interest is to identify SE genes from spatially resolved read count data. Suppose we have gene expression levels measured from *m* genes across *n* spatial spots on a target tissue. Below we focus on one gene and describe how to obtain a *p*-value for this gene. We refer to each spatial spot as a sample and denote the spatial coordinate as ***s***_*i*_ = (*s*_*i*1_, *s*_*i*2_), *i* = 1, …, *n*.

The SPARK algorithm models the expression count data of the focal gene across *n* spatial samples with a GLSM:

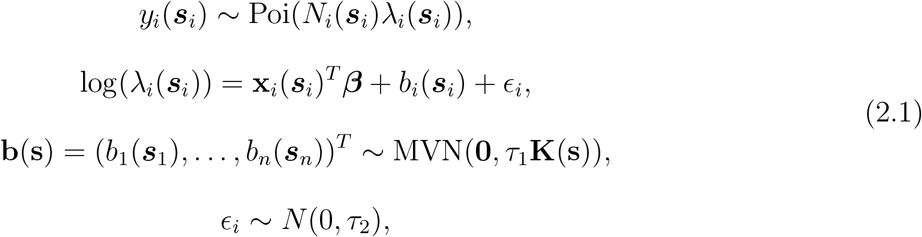

where *y_i_*(***s**_i_*) denotes the gene expression counts of the focused gene at the *i*th sample (spot); *N_i_*(***s**_i_*) denotes the total read counts of all genes for the *i*th sample; λ*_i_*(***s**_i_*) is an unknown Poisson rate parameter representing the underlying gene expression level of the focused gene at the *i*th sample. To explore the spatial pattern of the gene, log(λ*_i_*(***s**_i_*)) is modeled as a linear combination of three terms, where **x***_i_*(***s**_i_*) denotes a *k*-dimensional vector of explanatory variables for the *i*th sample (including one for the intercept and *k* – 1 observed explanatory variables that contain batch information, cell-cycle information, or other information that is important to adjust for during the analysis), ***β*** is a *k*-dimensional regression coefficient, *b_i_*(***s**_i_*) is a zero-mean, stationary Gaussian process modeling the spatial correlation pattern, and 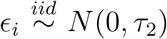 is the residual error independent of spatial locations. It is assumed that **b**(**s**) follows a multivariate normal distribution with mean **0** and covariance matrix *τ*_1_**K**(**s**), where *τ*_1_ is a scaling factor and **K**(**s**) is a kernel function of the spatial location **s** = (***s***_1_, …, ***s**_n_*)^*T*^, with the *ij*th element being **K**(***s**_i_*, ***s**_j_*).

Testing whether a gene shows spatial expression patterns is equivalent to testing the null hypothesis

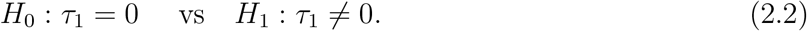

An approximate inference algorithm based on the penalized quasi-likelihood (PQL) algorithm (Breslow and Clayton, 1993) is developed to iteratively estimate ***β*** and *τ*_2_ under *H*_0_, enabling scalable analysis of tens of thousands of genes across hundreds of spatial locations. Then a score test statistic following a mixture of χ^2^ distributions under *H*_0_ is constructed based on the estimated parameters and the kernel function **K**(**s**) as in model (2.1). SPARK uses five Gaussian kernels and five cosine kernels with different pre-determined smoothness and periodicity parameters to match the true underlying spatial pattern displayed by the gene in focus. With each of the ten kernels, a *p*-value of the focal gene can be computed from corresponding score statistic. Finally, the *p*-values computed from the ten kernels are combined into a single *p*-value through the recently developed Cauchy combination rule (Liu *et al*., 2019; Liu and Xie, 2020) to obtain an overall *p*-value of the focal gene. Detailed calculation and implementation can be found in Sun *et al*. (2020).

After obtaining the *p*-values of *m* genes of our primary interest, we next use OrderShapeEM to incorporate auxiliary information into the multiple testing procedure. The Cauchy combination rule is utilized to integrate multiple auxiliary sequences. we provide detailed explanation in Section *Combination of multiple auxiliary sequences*.

### 2.2 Combination of multiple auxiliary sequences

In omics studies, auxiliary information could come from multiple sources. The original integrative analysis in OrderShapeEM is designed to have one sequence of auxiliary information. In this section, we propose a way to combine multiple sequences of auxiliary information to be used in OrderShapeEM. It is assumed that we have summary statistics, such as *p*-values from auxiliary sequences. We use *p*-values due to their generality, irrespective of underlying data type, statistical modeling and test statistics, among others. We require that *p*-values from auxiliary studies all have measurements on *m* genes. As usual, the *p*-values are assumed to follow a standard uniform distribution under the null hypothesis. Suppose we have *D* studies with respective *p*-values {*p_ji_*, *j* = 1, …, *m*; *i* = 1, …, *D*}. The auxiliary covariate for gene *j* can be formed by

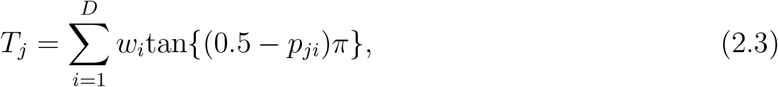

where the non-negative weights have to satisfy 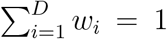 and can be used to improve informativeness of the auxiliary information. For instance, studies with larger sample size may have heavier weights. When *p_ji_* ~ *U*(·), tan{(0.5 – *p_ji_*)*π*}, *i* = 1, …, *D* follow standard Cauchy distributions. It has been shown that heavy tail outweighs correlation for Cauchy distributions (Pillai and Meng, 2016; Cohen *et al*., 2020; Liu and Xie, 2020). In Eq (2.3), the distribution of *T_j_* is standard Cauchy for any finite *D* and any dependence among *p_ji_*’s, *i* = 1, …, *D* when *p_ji_* ~ *U*(·). Define the covariate

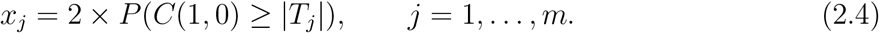

where *C*(1,0) stands for standard Cauchy distribution. Alternatively, we can obtain the covariate

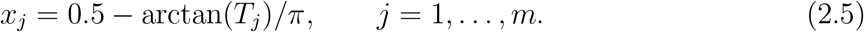

#### Algorithm 1: Identify SE genes from spatial transcriptomic data incorporating multi-omics auxiliary covariates

**Step 1. Obtain primary *p*-values from spatially resolved transcriptomic data with SPARK.** Given a primary spatial transcriptomic dataset containing read counts of *m* genes on *n* spatial spots. For each focal gene *j*, SPARK models its raw count data across *n* spots as a GLSM, and calculate the corresponding primary *p*-value via a score test with parameters inferred from model (2.1) under the null hypothesis *τ*_1_ = 0. Repeat the process for *m* genes, the primary *p*-values 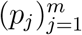 can be obtained.

**Step 2. Construct the auxiliary covariate sequence from multi-omics data.** Given *D* omics datasets related to the primary study, make appropriate gene-level statistical analysis on each dataset to obtain *D* auxiliary *p*-value sequences. If *D* > 1, match the *D* sequences by gene and combine them using the Cauchy combination rule, resulting in the auxiliary covariate sequence 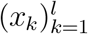.

**Step 3. Identify SE genes with OrderShapeEM.** Match 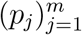 and 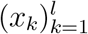 by genes. Without loss of generality, letting the matched pairs be 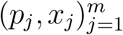, OrderShapeEM estimates 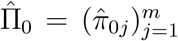 and 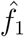 with a combination of PAVA and EM algorithm by introducing a monotone constraint between 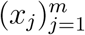 and 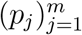. Get the order statistics 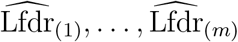. The SE genes are obtained by rejecting hypotheses corresponding *H*^(*i*)^ for 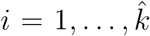 at a pre-specified FDR control level *α*, where 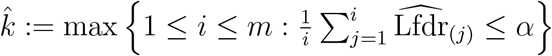.

Algorithm 1 summarizes our integrative analysis method that uses OrderShapeEM to detect SE genes from spatial transcriptomic data incorporating multi-omics auxiliary information, which is implemented in three steps. By integrating datasets from multiple studies, we reduce false discoveries as the probability to have large sporadic values across multiple studies is low. The method is robust to mis-specification of auxiliary information. Even if the covariate does not provide useful information, we still have asymptotic FDR control with comparable power to analysis using only the primary study dataset.

### 2.3 Multi-omics integrative analysis

In this section, we describe in detail the Lfdr-based multiple testing procedure incorporating the combined auxiliary information from multi-omics data. Suppose we have *m* primary hypotheses *H*_1_, …, *H_m_*, which test for the spatially expressed patterns of *m* genes. Denote *θ_j_* as the hidden true state indicator, where *θ_j_* = 1 indicates that the *j*th gene is a SE gene and *θ_j_* = 0 otherwise. Unlike classic multiple testing procedure that assumes exchangeability among *H*_1_, …, *H_m_*, we assume the probability that *θ_j_* = 0 varies across different genes. Specifically,

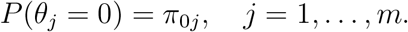

Conditional on *θ_j_*, we impose a two-group mixture model on the primary *p*-values (Efron, 2008):

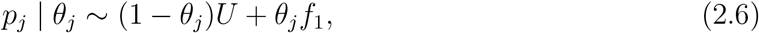

where *f*_1_ is the *p*-value density function for SE genes and *U* denotes standard uniform density for *p*-values under the null. Lfdr is the posterior probability that a gene is null given the corresponding *p*-value is equal to *p*. The Lfdr statistic is defined as

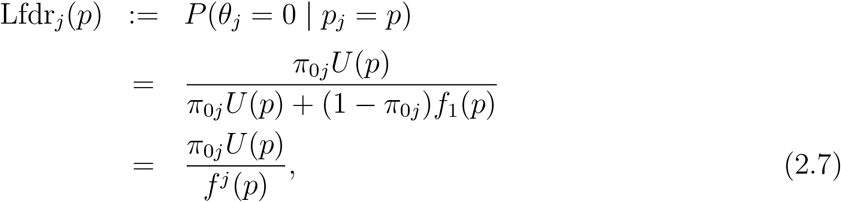

where

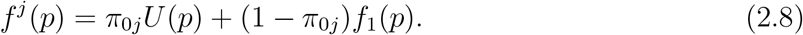

Under the monotone likelihood ratio assumption (Sun and Cai, 2007; Cao *et al*., 2013):

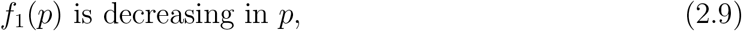

Lfdr_*j*_(*p*) is monotonically increasing in *p* (Cao *et al*., 2022). As a result, the rejection region can be represented as

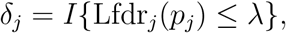

for a constant λ. We write the number of false rejections

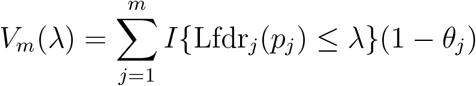

and the number of total rejections

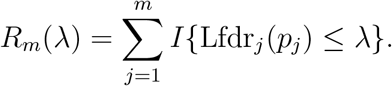

The FDR is defined as

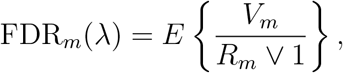

where *a* ∨ *b* = max{*a, b*}.

Note that

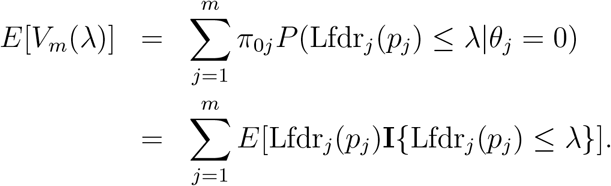

Consequently, an estimate of the FDR_*m*_(λ) is

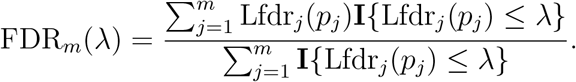

Define

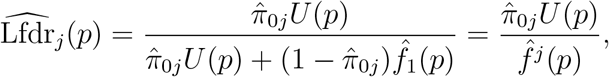

where 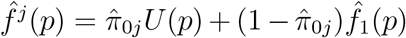. At a pre-specified FDR level *α*, a natural threshold is

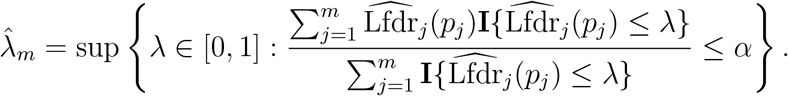

Reject the *j*th hypothesis if 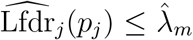. This is equivalent to the following step-up procedure that was originally proposed in Sun and Cai (2007). Let 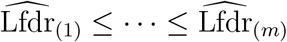 be the order statistics of 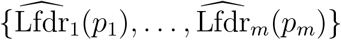 and denote by *H*^(1)^, …, *H*^(*m*)^ the corresponding ordered hypotheses. Define

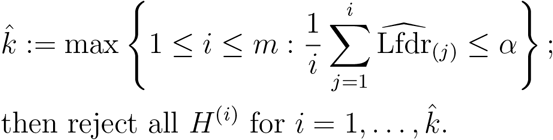

To implement this procedure, we need to estimate *m* unknown parameters *π*_0*j*_, *j* = 1, …, *m* and an unknown function *f*_1_(·). With a bivariate 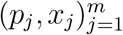, we introduce a monotone constraint on *π*_0*j*_, *j* = 1, …, *m* and *f*_1_(·) with respect to 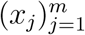 to estimate them by maximizing the log likelihood:

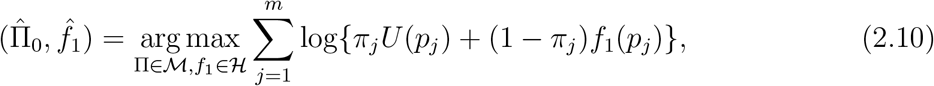

where 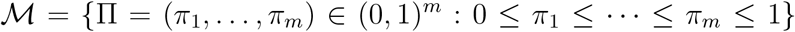 is a convex set, and 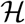 is the class of decreasing density functions. An illustrative example can be used to motivate the monotone constraint (see Supplementary Material Section A). With this constraint, the order of *x_j_* generates a ranked list of the primary hypotheses *H*_1_, …, *H_m_*.

Therefore, we can combine PAVA and EM to obtain estimation of *π*_0*j*_, *j* = 1, …, *m* and *f*_1_(·) incorporating the auxiliary covariates *x_j_*, *j* = 1, …, *m*.

- Step 1. Input the initial values 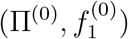.
- Step 2. **E-step:** Given 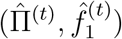, let

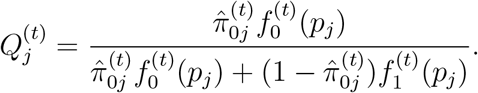
- Step 3. **M-step:** Given 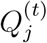, update (∏, *f*_1_) through

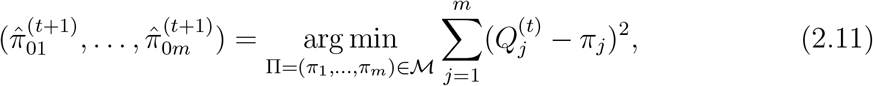

and

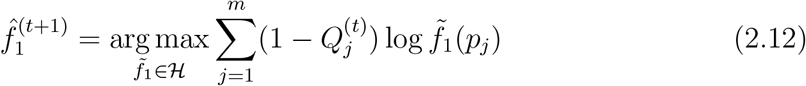
- Step 4. Repeat the E-step and M-step until the algorithm converges.

As EM algorithm is a hill-climbing algorithm, the above algorithm always converges. The minimization in Eq (2.11) and (2.12) can be accomplished through PAVA. If 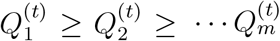, then the solution to Eq (2.11) is simply given by 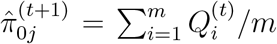 for all 1 ≤ *j* ≤ *m*. In other words, when the auxiliary ordering is mis-specified, the problem reduces to the case that prior probabilities of being null are exchangeable. Detailed derivation of this constrained optimization problem can be found in Cao *et al*. (2022).

## 3 Results

### 3.1 Realistic simulations

In this section, we investigate the performance of the proposed integrative analysis method through a series of realistic simulations. As benchmarks, in addition to the original SPARK algorithm that adopts the BY procedure (Benjamini and Yekutieli, 2001) for FDR control, we also implement the classic multiple testing procedure that does not incorporate any external information (BH) and another multiple testing procedure exploiting auxiliary information (SABHA). Implementation and detailed descriptions of these methods can be found in Supplementary Material Section A.

Following the simulation design in Sun *et al*. (2020), we generated gene expression data guided by the spatially resolved mouse olfactory bulb data (see Section *Mouse olfactory bulb data* for details) as well as parameters inferred from SPARK (*β* and *τ*_2_). The read counts for *m* = 10,000 genes were simulated. Without loss of generality, we set the first *ψ* (*ψ* = 7.5%, 10% or 15%) of the *m* genes as SE genes (non-null genes), and the remaining (1 – *ψ*)*m* genes as non-SE genes (null genes). In each setting, we simulated *y_i_*, *i* = 1, …, *n* for all *m* genes one by one, and then used SPARK to analyze the synthetic spatial transcriptomic data to obtain primary *p*-values for SE identification.

Differential analysis of RNA-seq data may yield informative auxiliary covariates that can be incorporated in the SE analysis. In our simulations, RNA-seq read counts for *m* = 10, 000 genes from two groups each consisting of 50 samples were generated. In accordance with the primary spatial data, *ψ* (*ψ* = 7.5%, 10% or 15%) of the *m* genes were simulated to be differentially expressed between the two groups (equally distributed between up- and down-regulation in group 2 compared to group 1). We explored the effect of the covariate at three different informative levels (i.e., “uninformative”, “weakly informative” and “strongly informative”) on the performance of the primary hypothesis testing. The uninformative covariates were generated from a standard uniform distribution *U*(0, 1). To generate a weakly informative covariate sequence, the differential expressed genes in the auxiliary dataset were randomly selected from all *m* = 10, 000 genes. For a strongly informative covariate sequence, we shrank the range of differentially expressed genes by randomly selecting from the first 1, 500, 2, 000 and 3, 000 genes, respectively. After obtaining the auxiliary RNA-seq dataset, the R package *DESeq2* v1.30.1 (Love *et al*., 2014) was applied to perform differential analysis, and the resulting *p*-values were utilized as auxiliary covariates for statistical analysis of SE genes. More simulation details can be found in Supplementary Material Section B.

For each simulation setting, 100 replicates were generated and analyzed by using different methods. FDR and power (true discovery rate) were calculated by averaging over the replications, where the power is defined as the expected value of the ratio of rejected SE genes over total number of true SE genes.

Fig. 2 presents FDR control and Fig. 3 shows power comparison of different methods. Across all simulation settings, the FDR is well controlled at the nominal level, which guarantees the validity. We observe that the BY method, which implements FDR control with arbitrary dependence, is most conservative, followed by BH, which does not take into account any external side information. With auxiliary information, we see power improvement, which is more pronounced with strongly informative auxiliary information for OrderShapeEM. In terms of computation, two tuning parameters are needed for SABHA, whereas OrderShapeEM is tuning parameter free and very easy to implement.

**Figure 2:**
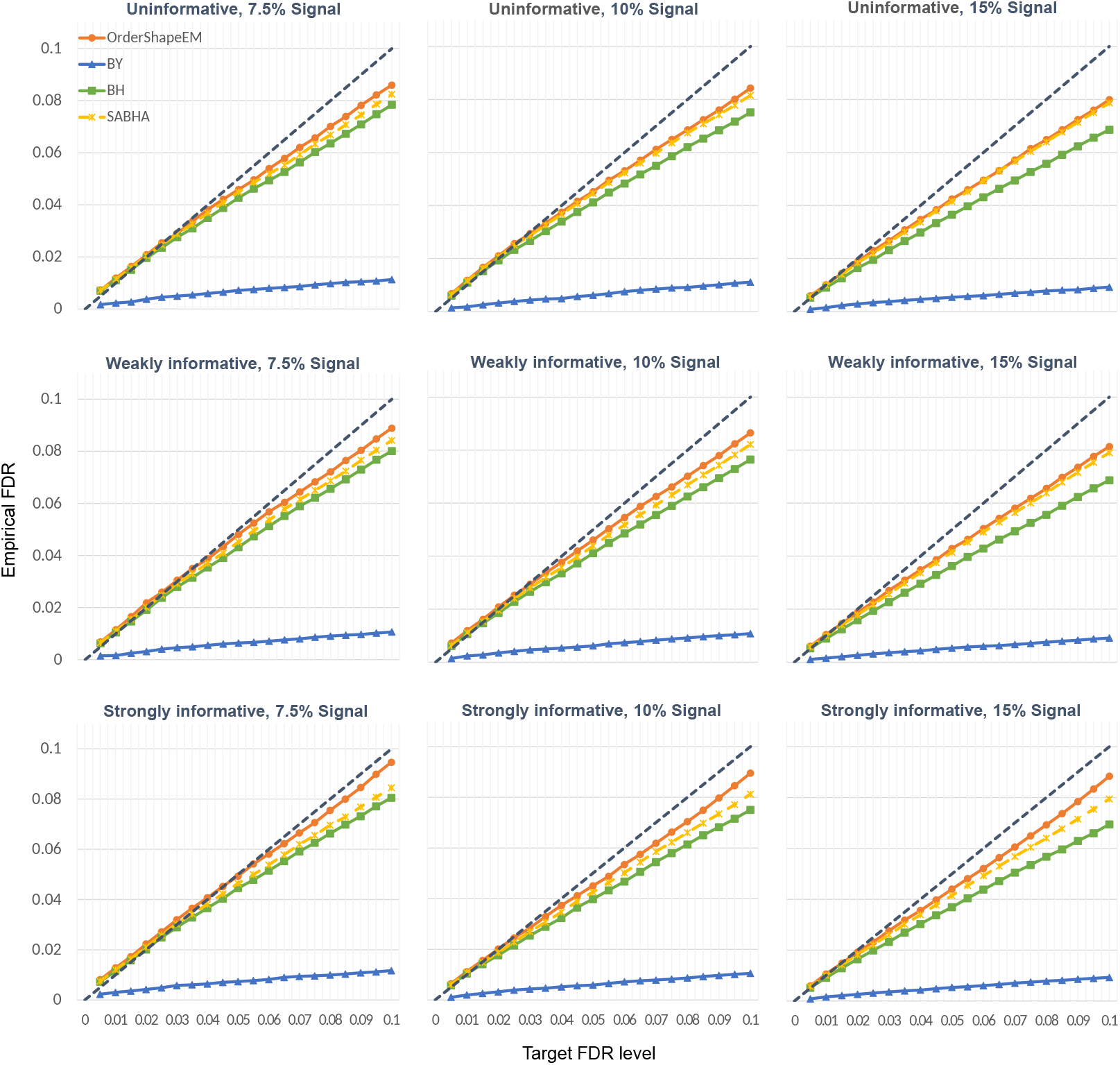
FDR comparison of different methods in the realistic simulations. Simulations were performed under three levels of non-null (SE genes) proportion (from left to right: 7.5%, 10%, 15%) and three levels of covariate informativeness (top: uninformative; middle: weakly informative; bottom: strongly informative).

**Figure 3:**
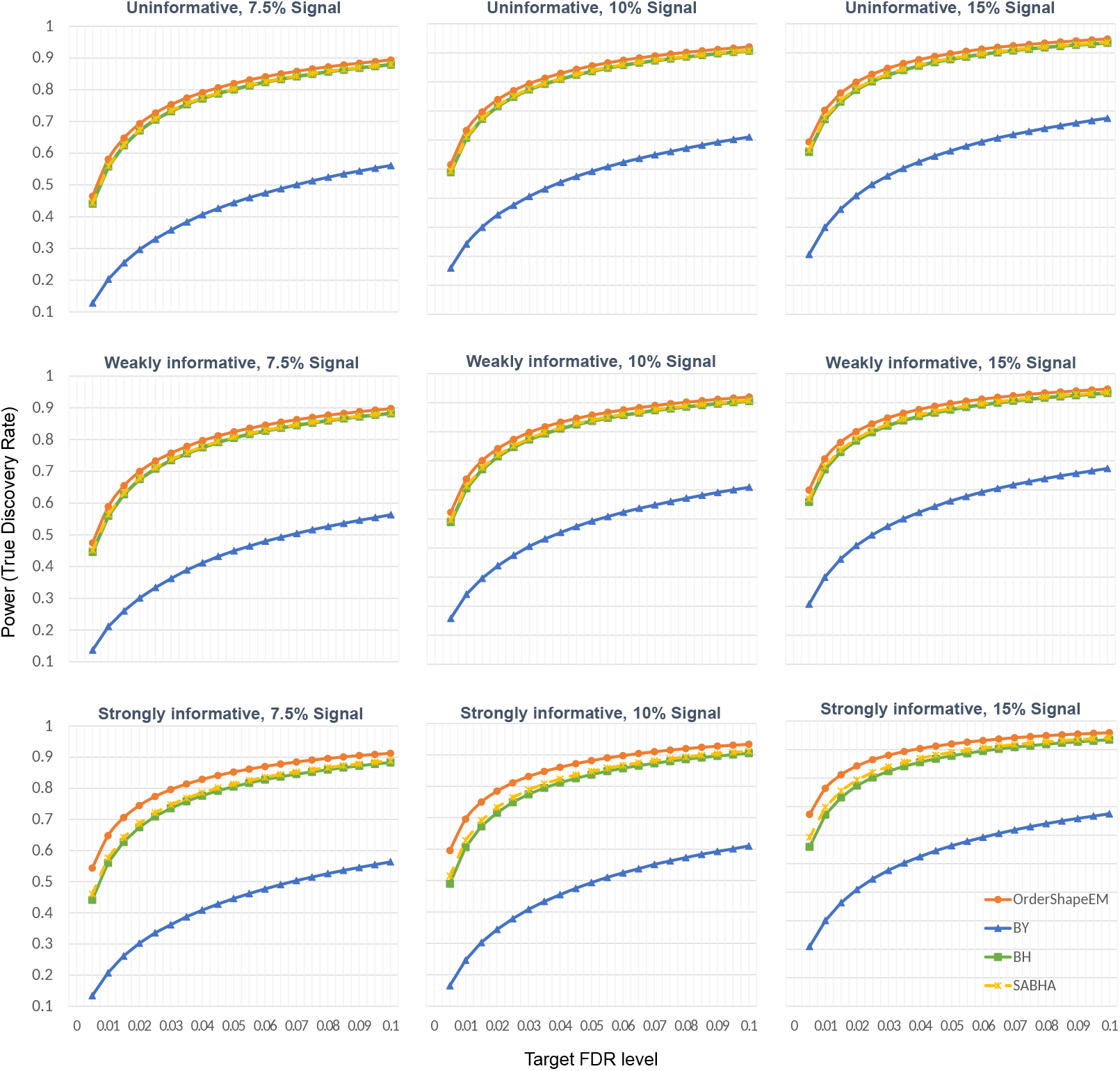
Power comparison of different methods in the realistic simulations. Simulations were performed under three levels of non-null (SE genes) proportion (from left to right: 7.5%, 10%, 15%) and three levels of covariate informativeness (top: uninformative; middle: weakly informative; bottom: strongly informative).

### 3.2 Mouse olfactory bulb data

We applied the proposed framework to analyze four spatial transcriptomic datasets (Stahl *et al*., 2016; Stickels *et al*., 2021): the mouse olfactory bulb data integrates scRNA-seq data, the mouse cerebellum data integrates scRNA-seq data, the human heart data integrates combined scRNA-seq and single nuclei RNA (snRNA)-seq data, and the human breast cancer data integrates The Cancer Genome Atlas (TCGA) RNA-seq data from the Breast Cancer program (BRCA), TCGA RNA-seq data from the Thyroid Cancer program (THCA) and GWAS summary data from the Breast Cancer Association Consortium (BCAC) for women of European descent. The data acquisition details can be found in Supplementary Material Section C.

First, we implemented the proposed integrative analysis framework on the mouse olfactory bulb data consisting of 11, 274 genes measured on 260 spots (Stahl *et al*., 2016). The SPARK algorithm was applied for SE analysis to obtain the primary *p*-values. We used a publicly available scRNA-seq dataset consisting of 27, 998 genes measured on 20, 636 mouse olfactory cells (Zeisel *et al*., 2018) as the auxiliary data. Differential expression analysis was performed on this dataset to find marker genes for each cell-type cluster compared to all the remaining cell types, yielding an auxiliary *p*-value sequence containing 8, 084 genes. We took the 7, 026 pairs of *p*-values corresponding to the intersection of genes from the two studies as input to the OrderShapeEM algorithm.

At an FDR level 0.05, the BY procedure, which is adopted by SPARK, identified 739 SE genes. OrderShapeEM detected 1,430 SE genes, including all SE genes identified by SPARK and improved the statistical power by 94%. To assess the quality of the SE genes detected by OrderShapeEM, we clustered the genes into three different groups with the R package *amap* v0.8-18 and categorized the gene expression level of the cluster centers into three distinct spatial expression patterns (Fig. 4(A) and Supplementary Material Fig. S1(A)): Pattern I corresponds to the mitral cell layer, Pattern II corresponds to the granular cell layer and Pattern III corresponds to the glomerular cell layer. We also applied BH and SABHA on the same dataset. Fig. 4(B) shows the number of SE genes identified by each method and the overlaps of different methods using the R package *UpSetR* v1.4.0 (Conway *et al*., 2017; Lex *et al*., 2014). We can see that incorporating auxiliary information noticeably improved the detection of SE genes.

**Figure 4:**
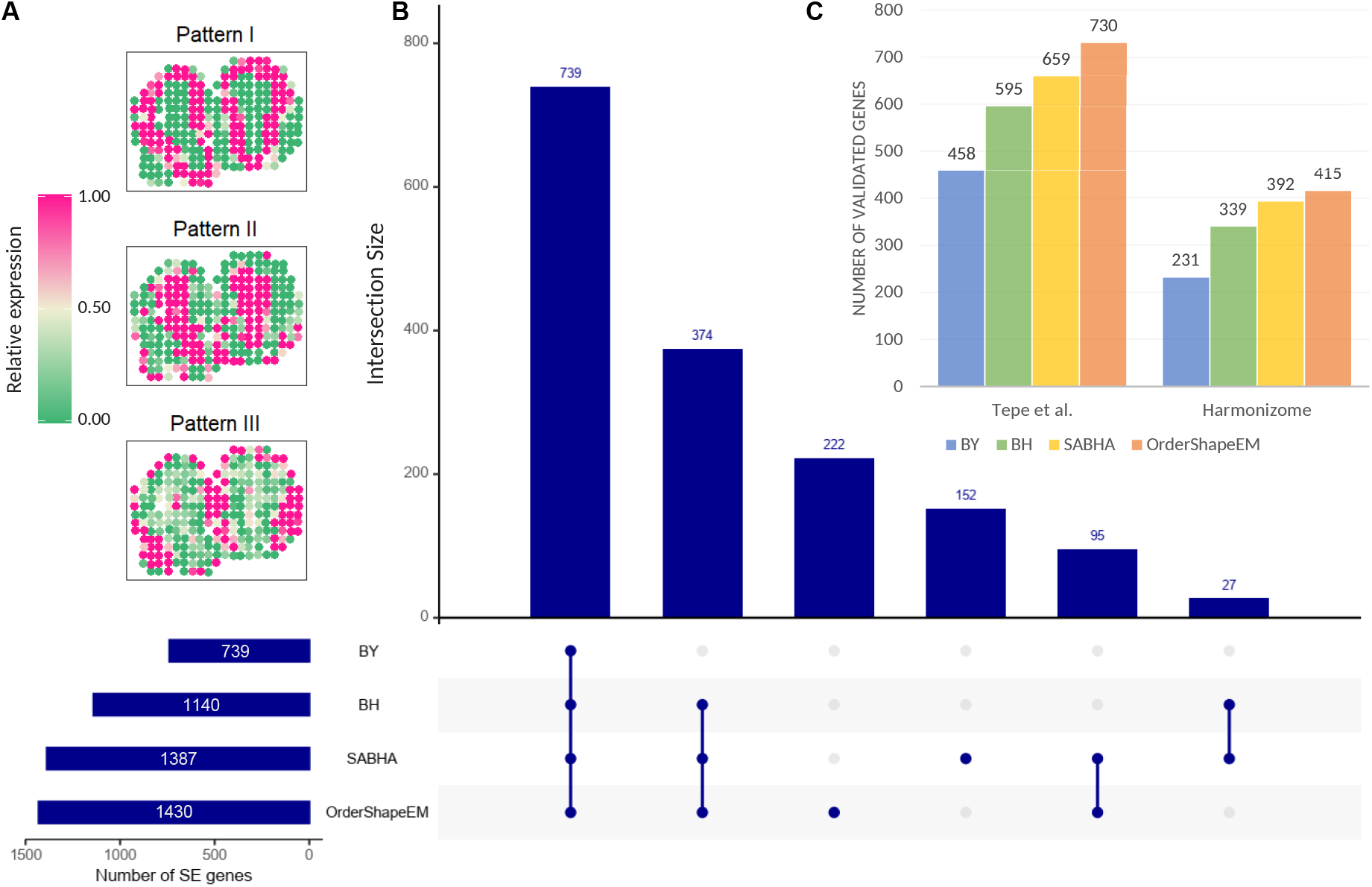
Analysis and validation results of the mouse olfactory bulb data. (A) Three clusters of distinct spatial expression patterns summarized on the basis of 1,430 SE genes identified by OrderShapeEM. The color represents relative gene expression levels: pink represents high expression regions and green indicates low expression regions. (B) Diagram shows the number of SE genes identified by different methods at a target FDR level *α* = 0.05 and their resulting discovery set intersections in a matrix layout. The horizontal bars show the total number of SE genes identified by each method; the vertical bars plot the number of SE genes in different intersection sets. The blue solid circle lights up when the SE genes identified by corresponding method is included in the intersection set. (C) Barplot displays the number of SE genes identified by each method that were validated in two reference gene lists: one from Tepe *et al*. (2018) and the other from the Harmonizome database.

We downloaded three reference gene lists from related studies published in public databases to further validate the SE genes identified by different methods. These genes exhibit spatial heterogeneity in various ways. First, we compared our findings to the highlighted SE genes in the mouse olfactory system identified by the original spatial transcriptomic study (Stahl *et al*., 2016). SPARK detected 7 of 10 such highlighted genes, BH, SABHA and OrderShapeEM identified one additional gene in the list, namely *Sv2b*, which plays an important role in the control of regulated secretion in neural cells (Janz *et al*., 1999). The second reference gene list is composed of 2, 030 cell-type-specific marker genes identified in a recent scRNA-seq study which reveals cellular heterogeneity in mouse olfactory bulb (Tepe *et al*., 2018). As shown in the left panel of Fig. 4(C), 458 of the 739 SE genes identified by SPARK were discovered in the reference gene list. In addition to the overlapping genes, of the 691 SE genes uniquely identified by OrderShapeEM, approximately 39% (272 SE genes) were validated in the list. Third, a gene list consisting of 3, 485 genes related to the three main mouse olfactory bulb layers (mitral, granular and glomerular) with high or low expression levels relative to other tissues was obtained from the Harmonizome database (Rouillard *et al*., 2016). The right panel of Fig. 4(C) exhibits the validation results: of all the 739 SPARK discoveries, 31% (231 SE genes) were validated in the Harmonizome gene list, whereas approximately 27% (184 SE genes) of the additional 691 SE genes uniquely identified by OrderShapeEM were also found in the gene list.

Finally, we performed Gene Ontology (GO) enrichment analysis on SE genes detected by SPARK and OrderShapeEM to gain biological insights. At a target FDR level of 0.05, 1, 557 GO terms were enriched in the 1,430 SE genes identified by OrderShapeEM, while 1, 281 GO terms were enriched in the SE genes identified by SPARK. Despite the overlaps (1, 133 GO terms), OrderShapeEM identified additional GO terms that are directly related to the neuronal synaptic organization and olfactory signaling pathways. For example, the GO terms only enriched by OrderShapeEM (GO:0007187 and GO:0007188) modulate G protein-coupled receptor signaling pathway, which plays a critical role in olfactory signal transduction, neuronal morphology and axon guidance of mouse (Ebrahimi and Chess, 1998). This demonstrates the advantage of incorporating external information in the analysis of spatial transcriptomic data.

### 3.3 Mouse cerebellum data

Second, we examined the application of the proposed integrative analysis framework to a mouse cerebellum dataset consisting of 523 genes measured on 11, 625 beads through Slide-seqV2 (Stickels *et al*., 2021). The dataset was obtained by filtering out genes that were expressed in less than 10% of the spatial locations and selecting spatial locations with at least ten total read counts for computational reasons. With SPARK, we obtained the primary *p*-values of these 523 genes. We then downloaded a public scRNA-seq dataset of mouse cerebellum cells (Zeisel *et al*., 2018) and performed differential expression analysis between different cell-types to obtain an auxiliary *p*-value sequence for 6, 666 genes. By taking intersection of genes in the primary and auxiliary study, we used 502 *p*-value pairs as input to OrderShapeEM.

As shown in Fig. 5(A), SPARK identified 315 SE genes at the target FDR level of 0.05, all of which were also discovered by other three methods (i.e., BH, SABHA and OrderShapeEM), and all 366 SE genes identified by BH were detected by SABHA and OrderShapeEM. By incorporating auxiliary information provided by the scRNA-seq data of mouse cerebellum, OrderShapeEM identified 459 SE genes, improving the power of original SPARK by 46%. To evaluate additional discoveries made by OrderShapeEM, we clustered the 459 SE genes identified into two different groups by using the R package *amap* v0.8-18 and categorized the gene expression level of the two clusters into two distinct spatial expression patterns: Pattern I corresponds to the granular cell layer and Pattern II displays multiple layers including the purkinje layer. From Fig. 5(B)(C) and Supplementary Material Fig. S2(A), we see that the clustered spatial expression patterns are consistent with the coronal section of mouse cerebellum in the Allen Brain Atlas.

**Figure 5:**
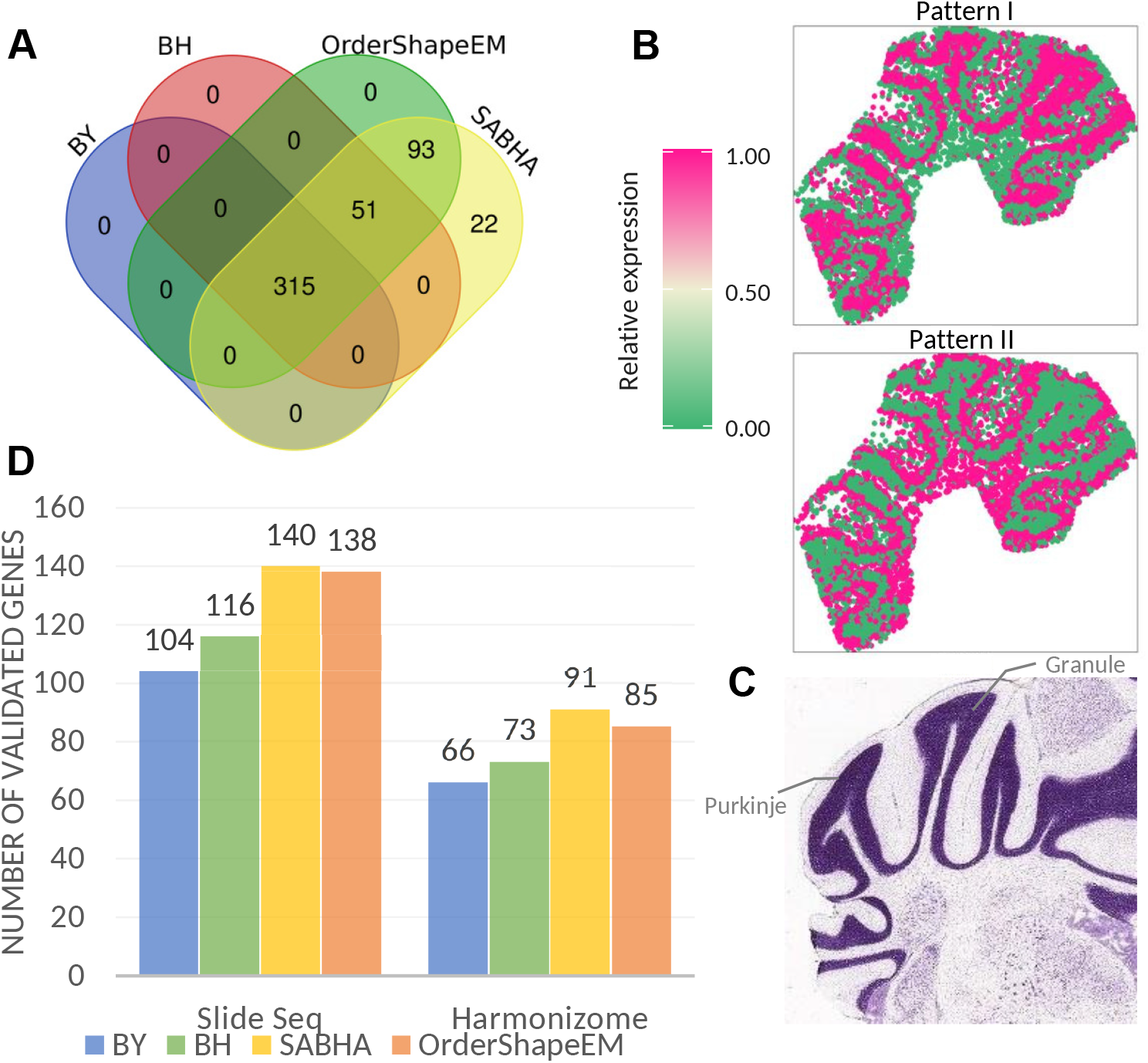
Analysis and validation results of the mouse cerebellum data. (A) Venn diagram shows the number of SE genes identified by different methods at a target FDR level *α* = 0.05 and their resulting discovery set intersections. (B) Two distinct spatial expression patterns summarized based on the 459 SE genes identified by OrderShapeEM. (C) Coronal section of mouse cerebellum obtained from the Allen Brain Atlas. (D) Barplot displaying the number of SE genes identified by each method that were validated in two reference gene lists: one from the Slide-seq study and the other from the Harmonizome database.

We then provide additional evidence to validate the discoveries based on two reference gene lists downloaded from previous related studies. First, we compared our results to 669 spatially relevant genes identified in the purkinje layer of mouse cerebellum in the original Slide-seq study (Rodriques *et al*., 2019). As shown in the left panel of Fig. 5(D), SPARK detected 104 of these highlighted genes. In addition to the overlapping genes with SPARK, OrderShapeEM identified additional 34 SE genes comparable to additional 36 genes identified by SABHA. Second, a reference gene list composed of 1, 867 genes with high or low expression in mouse cerebellum relative to other tissues was obtained from the Harmonizome database (Rouillard *et al*., 2016). As can be seen in the right panel of Fig. 5(D), 21% (66) of the 315 SE genes identified by SPARK were validated in the reference list. Moreover, additional 19 SE genes identified by OrderShapeEM were found in the same list. These results show that the proposed integrative analysis improves the power, and additional findings have biological relevance. We did not do GO enrichment analysis since a relatively small number of SE genes were identified.

### 3.4 Human breast cancer data

We also applied the proposed integrative analysis to a human breast cancer dataset comprised of 5, 262 genes on 250 spots (Stahl *et al*., 2016). We used SPARK to obtain the primary *p*-values from the spatial transcriptomic data. Next we downloaded three datasets related to human breast cancer: TCGA BRCA RNA-seq data consisting of 55, 112 genes, TCGA THCA RNA-seq data consisting of 39, 788 genes, and GWAS summary data from BACA for women of European ancestry consisting of 11, 089, 342 SNPs (Michailidou *et al*., 2017). We conducted differential expression analyses on the TCGA RNA-seq datasets and made gene-based association analysis on the GWAS summary statistics. These three auxiliary datasets can yield seven different types of auxiliary *p*-value sequences through free combinations, which are denoted as Case 1-7, respectively (Supplementary Material Table S1).

We took the intersection of three auxiliary datasets and the primary dataset, obtaining 4, 801 genes that were present in all studies. Next the *p*-value pairs of the common genes in each case were used as input to OrderShapeEM. We observe that the 268 SE genes identified by SPARK were also discovered by other methods (i.e., BH, SABHA and OrderShapeEM). BH discovered 498 SE genes with overwhelming overlaps with genes discovered by SABHA and OrderShapeEM. Fig. 6(B) presents the analysis results of different methods with different combination of auxiliary datasets for OrderShapeEM (Case 1-7) (Conway *et al*., 2017; Lex *et al*., 2014). When incorporating all three datasets as auxiliary covariates, 571 SE genes were detected by SABHA, whereas 906 SE genes were discovered by OrderShapeEM. We noticed that the more auxiliary datasets OrderShapeEM integrated the more discoveries were made. This reflected the strength of information borrowing across different studies. We further clustered the 906 SE genes identified by OrderShapeEM in Case 7 into two different groups and categorized the gene expression level of the cluster centers into two distinct spatial expression patterns (Fig. 6(A) and Supplementary Material Fig. S3(A)(B)). Pattern I as shown in Fig. 6(A) corresponds to the possible tumor region in the hematoxylin and eosin staining obtained from Sun *et al*. (2020).

**Figure 6:**
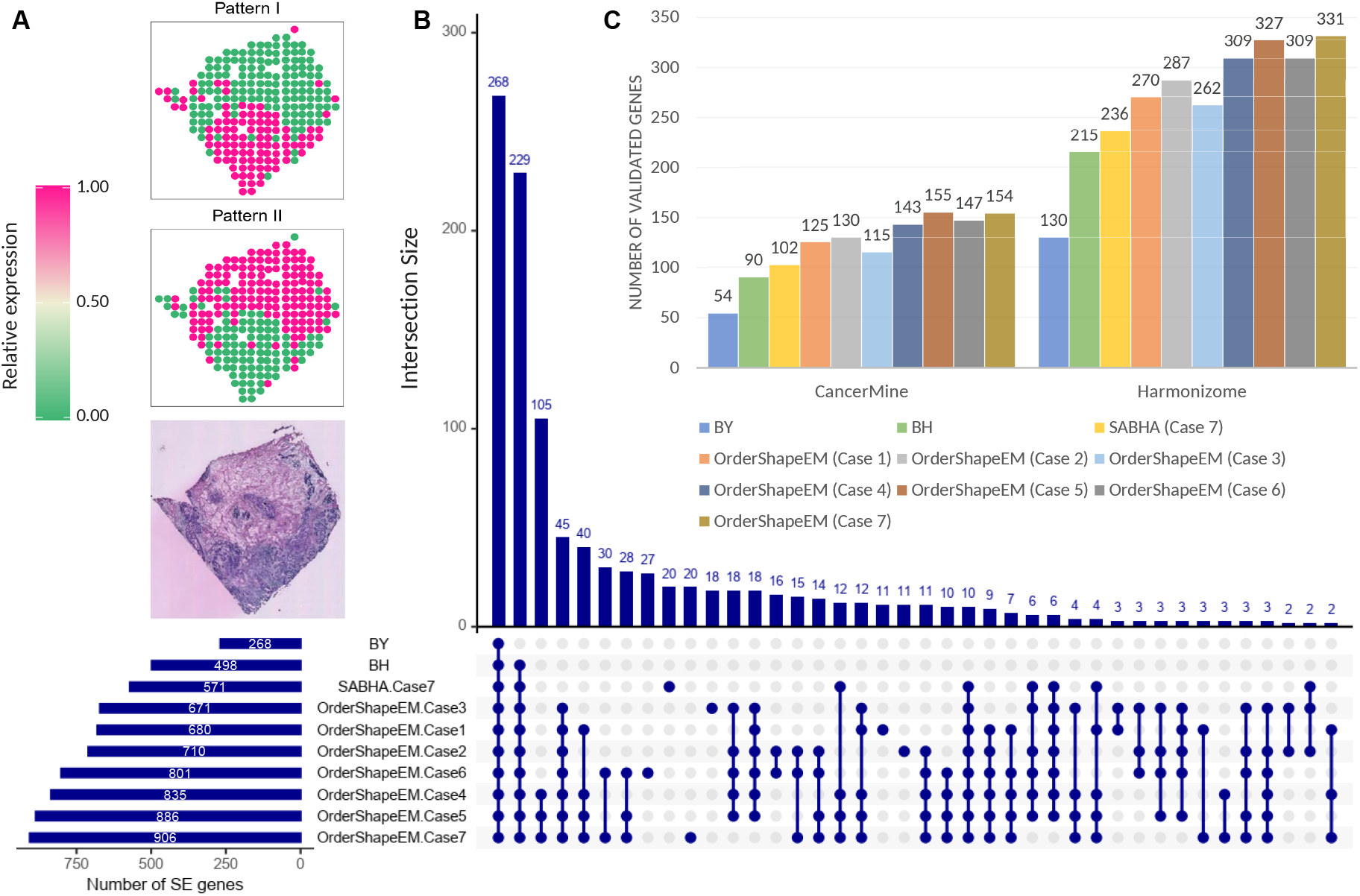
Analysis and validation results of the human breast cancer data. (A) Two cluster of spatial expression patterns based on the 906 SE genes identified by OrderShapeEM. The color represents relative gene expression levels: pink represents high expression regions and green indicates low expression regions. The hematoxylin and eosin staining are obtained from Sun *et al*. (2020) and shown at the bottom, with dark staining representing the possible tumor region. (B) Diagram shows the number of SE genes identified by different methods at a target FDR level *α* = 0.05 and their resulting discovery set intersections in a matrix layout. The horizontal bars show the total number of SE genes identified by each method (in different cases); the vertical bars plot the number of SE genes in different intersection sets. The blue solid circle lights up when the SE genes identified by corresponding method is included in the intersection set. (C) Barplot displays the number of SE genes identified by each method that were validated in two reference gene lists: one from the CancerMine database and the other from the Harmonizome database.

We further validated the quality of additional discoveries with three reference gene lists which provided valuable insights into SE analysis by showing differential expression patterns in human breast cancer tissues under different conditions. First, we examined the 14 marker genes which were highlighted to show spatial expression patterns in human breast cancer biopsies in Stahl *et al*. (2016). For all 7 cases, BH, SABHA and OrderShapeEM detected 13 of the 14 marker genes, whereas SPARK identified 10 of them. The three genes missed by SPARK (*MUCL1, GAS6 and AREG*) have been widely studied and well recognized to be related to breast cancer. Second, we obtained a list of 1, 642 genes relevant to breast cancer from the CancerMine database (Lever *et al*., 2019). As shown in the left panel of Fig. 6(C), 100 SE genes uniquely identified by OrderShapeEM were in the CancerMine list in Case 7, whereas 54 of the 268 SPARK discoveries were in the same list. Third, we collected another reference gene list from the Harmonizome database (Rouillard *et al*., 2016), including 3,505 breast cancer related genes from six different studies: OMIM gene-disease associations, PhosphoSitePlus phosphosite-disease associations, DISEASES text-mining gene-disease association evidence scores, GAD gene-disease associations, GWAS catalog SNP-phenotype associations, and GWASdb SNP-disease associations. The right panel of Fig. 6(C) displays the validation result, OrderShapeEM has more validated SE genes in the Harmonizome list.

Then we performed GO enrichment analysis on the SE genes identified by SPARK and OrderShapeEM. At a target FDR level of 0.05, SPARK enriched 553 GO terms, whereas OrderShapeEM identified 650 GO terms in Case 2 analysis. In addition to the 430 overlapped GO terms, OrderShapeEM generated many enrichment for biological processes related to cancer development. For example, the GO term of leukocyte activation involved in immune response (GO:0002366) and five other GO terms related to immune response were only enriched by OrderShapeEM. The vital role of the immune system in breast cancer initiation and development were recognized by many studies (Emens, 2012; Gonzalez *et al*., 2018).

### 3.5 Human heart data

We finally applied the integrative analysis framework to a 10X Visium spatial expression data from the human heart tissue. The filtered dataset contains 20, 904 genes measured on 4, 247 spots. A public available combined scRNA-seq and snRNA-seq dataset of 485K cardiac cells (Litviňuková *et al*., 2020) was used as auxiliary data. Differential expression analysis between cell-types produced an auxiliary *p*-value sequence for 4, 088 genes, of which 3, 848 genes were also in the primary study. We performed SPARK analysis on the spatial expression data of these 3, 848 overlapping genes on 4, 247 spots and obtained the corresponding primary *p*-values.

At a target FDR level of 0.05, OrderShapeEM detected all SE genes discovered by the other three methods (Fig. 7(A)). Compared to the 330 SE genes identified by SPARK (BY), OrderShapeEM identified additional 494 SE genes. To evaluate the spatially expressed patterns of the SE genes detected by OrderShapeEM, we performed clustering on the 824 genes and obtained two spatial expression patterns. Fig. 7(C) plots the distribution of cells in each cluster by using the UMAP (R package *umap* v0.2.8.0) to reduce the dimensionality into two dimensions. We also computed the Moran’s *I* statistic (Moran, 1950), a commonly used metric for quantifying spatial autocorrelation, to demonstrate the spatial variation of the OrderShapeEM discoveries (Fig. 7(D)). It can be seen that the 494 SE genes uniquely identified by OrderShapeEM have higher Moran’s *I* statistics than those not identified by OrderShapeEM.

**Figure 7:**
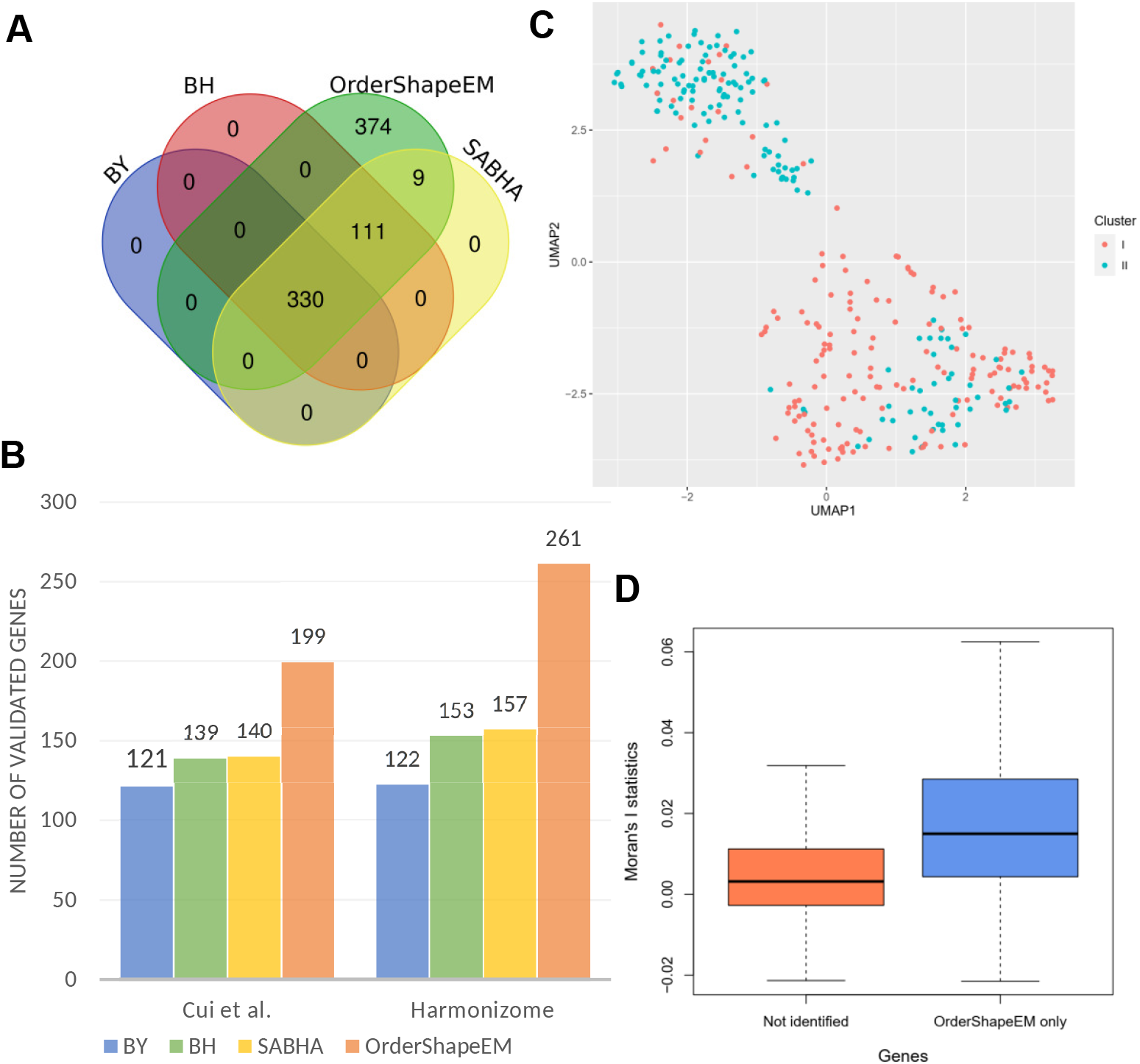
Analysis and validation results of the human heart data. (A) Venn diagram shows the number of SE genes identified by different methods at a target FDR level *α* = 0.05 and results intersections. (B) Bar charts displays the number of SE genes identified by each method that were validated in two reference gene lists: one from (Cui *et al*., 2019) and the other from the Harmonizome database. (C) Scatter plot of 824 SE genes identified by OrderShapeEM. We first use the UMAP method to reduce the dimensionality to two dimensions. We then use the two-factor cell labels obtained from hierarchical agglomerative clustering to visualize the distribution of cells in each cluster. (D) Boxplot displays Moran’s *I* statistics for the 494 SE genes uniquely identified by OrderShapeEM and genes not identified by OrderShapeEM.

We provided two additional lines of evidence to validate the OrderShapeEM discoveries. First, we compared our results to a list of 1,428 differentially expressed genes published in a previous scRNA-seq analysis of the human heart (Cui *et al*., 2019). As shown in the left panel of Fig. 7(B), SPARK detected 121 of the highlighted genes. OrderShapeEM detected additional 78 SE genes in the reference gene list. Second, a list of 6, 723 genes with high or low expression in heart relative to other cell types and tissues from the BioGPS human cell type and tissue gene expression profiles dataset was obtained from the Harmonizome database (Rouillard *et al*., 2016). As can be seen from the right panel of Fig. 7(B), 122 of the 330 SE genes identified by SPARK were validated in the Harmonizome list. In addition, 261 of the 824 SE genes detected by OrderShapeEM were also validated in the same list.

We also performed GO enrichment analysis on the 330 SE genes identified by SPARK and 824 SE genes detected by OrderShapeEM. At a target FDR level of 0.05, SPARK and OrderShapeEM enriched 384 and 499 GO terms, respectively. Of the 203 significant GO terms uniquely identified by OrderShapeEM, many were enriched for biological processes related to cardiac function, such as the regulation of heart rate (GO:0002027), cell growth, differentiation and communication involved in cardiac (e.g., GO:0061049, GO:0055007, GO:0086064 and GO:0003161).

## 4 Discussion

Identifying genes that display spatial expression patterns is an important first step toward characterizing the spatial transcriptomic landscape in tissues. In this paper, we extend the integrative analysis method OrderShapeEM with the Cauchy combination rule to allow covariate-assisted multiple hypothesis testing exploiting auxiliary data from multiple sources. We apply the extended OrderShapeEM method to improve statistical power of identifying SE genes from spatial transcriptomic data by incorporating multi-omics external auxiliary covariates, while maintaining FDR control. We use SPARK, a parametric modeling method, to model the primary spatial transcriptomics data, which can be vulnerable to model mis-specification. As an alternative, non-parametric SPARK-X is robust, computationally efficient and especially suitable when count data are sparse and the number of spatial spots is large (Zhu *et al*., 2021). The trade off is that SPARK-X usually has lower power compared with SPARK. We also implemented SPARK-X in the human heart data, the analysis results is available in the Supplementary Material Section D.4.

The integrative analysis framework we propose is very general. For instance, the primary study may have genes with spots on three dimensions. Our method is robust. We have improved power with informative auxiliary information. When auxiliary information is weakly informative or uninformative, we have comparable power with methods that do not incorporate auxiliary information. Our method is computationally scalable with the built-in PAVA and EM algorithms. Our method does not involve any tuning parameters, which allows easy implementation. We only require *p*-values from the primary study and auxiliary studies. This is very flexible as *p*-values can be obtained from parametric methods or non-parametric methods, such as permutation. Simulation studies and data analysis evidence that our method can reliably detect signals of biological relevance.

In this paper, we use spatially resolved transcriptomic data as an example, auxiliary information from scRNA-seq data, bulk RNA-seq data and GWAS summary data are considered. Our framework is applicable for other types of multi-omics data, such as ATAC-seq and CITE-seq after pre-processing. For example, the single cell ATAC-seq (scATAC-seq) data can be used to calculate *p*-values for each accessible peak by performing a differential accessibility test between clusters of cells. To incorporate such auxiliary data into our framework by gene, we can annotate each peak with the closest gene and use the corresponding *p*-values as the auxiliary covariate. One caveat is that our integrative analysis is a marginal based approach and does not incorporate dependence information such as linkage disequilibrium into account. We require that the primary and auxiliary data have overlapped genes, which may be difficult to obtain.

To ensure robust inference, we use the Cauchy combination rule to integrate multiple omics datasets into one sequence of *p*-values. The Cauchy combination rule guarantees that *p*-value distribution under the null hypothesis is standard uniform regardless of the dependence among auxiliary datasets (Pillai and Meng, 2016). Fisher’s combination rule for independent *p*-values is another option to integrate multiple omics datasets. The resulting *p*-values after the combination may no longer follow the standard uniform distribution under the null when the input *p*-values from multiple studies are correlated with each other (Fisher, 1925). Additional research is warranted to investigate the pros and cons of using different combination rules.

In the case that test statistic of SE genes in the primary analysis is in the opposite direction to that in auxiliary information, our method regards such auxiliary information to be informative. From *p*-values alone, we do not have the sign information of the test statistic. We do not require sign consistency between the primary and auxiliary studies. Sign consistency is an important criterion in replicability studies (Zhao *et al*., 2020).

## Data and codes availability

This study used four publicly available spatial transcriptomic datasets. The mouse olfactory bulb and human breast cancer data were downloaded from the Spatial Research at https://www.spatialresearch.org/resources-published-datasets/doi-10-1126science-aaf2403; the mouse cerebellum data is available at the Broad Institute’s single-cell repository (https://singlecell.broadinstitute.org/single_cell/) with ID SCP948; and the human heart data was collected from the 10X Visium spatial gene-expression repository (https://support.10xgenomics.com/spatial-gene-expression/datasets/1.1.0/V1_Human_Heart). The auxiliary scRNA-seq data used for mouse olfactory bulb and mouse cerebellum data are publicly available at http://mousebrain.org/adolescent/tissues.html. The auxiliary GWAS data for human breast cancer is available at the BCAC (http://bcac.ccge.medschl.cam.ac.uk/bcacdata/oncoarray). The auxiliary combined scRNA-seq and snRNA-seq data of cardiac cells is available at https://cells.ucsc.edu/?ds=heart-cell-atlas.

The R source code and scripts used in our analyses as well as underlying raw data are freely available at GitHub https://github.com/YanLi15/OrderShapeEM_SE. The implementation of the original SPARK and OrderShapeEM are available at GitHub https://github.com/xzhoulab/SPARK and https://github.com/jchen1981/OrderShapeEM, respectively.

