## Supplementary Material for "Statistical analysis of spatially resolved transcriptomic data by incorporating multi-omics auxiliary information"

Yan Li[1], Xiang Zhou[2] and Hongyuan Cao[3]

#### A Methods comparison overview

In addition to the original SPARK algorithm that adopts the BY procedure (Benjamini and Yekutieli , 2001) for FDR control, we also implement the classic multiple testing procedure that does not incorporate any external information (BH) and one modern multiple testing procedure exploiting auxiliary information (SABHA) as comparisons.

Given paired primary $p$-values and auxiliary covariates $(p_j, x_j)_{j=1}^m$ for hypotheses $H_j, j = 1, \ldots, m$, we implement the following multiple testing procedures in $R$ 4.0.5 to control FDR at target level $\alpha \in [0, 1]$.

- BY: Benjamini-Yekutieli procedure (Benjamini and Yekutieli , 2001) utilized by original SPARK, implemented by *p.adjust* function with *method="BY"*;
- BH: Benjamini-Hochberg procedure (Benjamin and Hochberg , 1995), implemented by *p.adjust* function with *method="BH"*;

[1] School of Mathematics, Jilin University, Changchun, Jilin 130012, China
[2] Department of Biostatistics, University of Michigan, Ann Arbor, MI 48109, U.S.A.
[3] Department of Statistics, Florida State University, Tallahassee, FL 32306, U.S.A.

- SABHA: Structure adaptive BH procedure (Li and Barber , 2019), implemented with parameters $\tau = 0.5, \epsilon = 0.1$;
- OrderShapeEM: Lfdr-based step-up procedure imposing a monotonic relationship between auxiliary sequence and $p$-values from the primary study (Cao *et al.* , 2022), implemented by *OrderShapeEM* function in the R package *OrderShapeEM* v1.0;

## A.1 BH

BH (Benjamin and Hochberg , 1995) is the most popular multiple testing procedure, which conservatively controls the FDR for $m$ independent or positively correlated tests. In our setups, the BH procedure proceeds as follows:

- Step 1. Let $p_{(1)} \leq p_{(2)} \leq \cdots \leq p_{(m)}$ be the ordered primary $p$-values, and denote by $H_{(j)}$ the null hypothesis corresponding to $p_{(j)}$;
- Step 2. Let $\hat{k}$ be the largest $j$ such that $p_{(j)} \leq \frac{j}{m}\alpha$, i.e.,

$$\hat{k} = \max\{j \geq 1 : p_{(j)} \leq \frac{j}{m}\alpha\},$$

  and $\hat{k} = 0$ if the set is empty;
- Step 3. Reject all $H_{(j)}$ for $j = 1, \ldots, \hat{k}$.

## A.2 BY

BY (Benjamini and Yekutieli , 2001) controls the FDR under arbitrary dependence. It is much more conservative than BH. After Obtaining primary $p$-values $p_j, j = 1, \ldots, m$, BY works as follows:

- Step 1. Let $p_{(1)} \leq p_{(2)} \leq \cdots \leq p_{(m)}$ be the ordered primary $p$-values, and denote by $H_{(j)}$ the null hypothesis corresponding to $p_{(j)}$;
- Step 2. Let $\hat{k}$ be the largest $j$ such that $p_{(j)} \leq \frac{j}{m \cdot c(m)}\alpha$;

  (a) If the tests are independent or positively correlated, $c(m) = 1$, which reduces to BH;

  (b) Under arbitrary dependence, $c(m)$ is the harmonic number: $c(m) = \sum_{j=1}^{m} \frac{1}{j} = \log m + \mathcal{O}(1)$;
- Step 3. Reject all $H_{(j)}$ for $j = 1, \ldots, \hat{k}$.

#### A.3 SABHA

SABHA (Li and Barber , 2019) incorporates auxiliary information about any pre-determined type of signal structure to reweigh the primary $p$-values in a covariate-adaptive way. Given a target FDR level $\alpha$ and a threshold $\tau \in [0, 1]$, SABHA works as follows:

- Step 1. Screen the primary $p$-values $p_j, j = 1, \ldots, m$ with threshold $\tau$, leading to censored $p$-values $p_j \mathbf{1}\{p_j > \tau\}$;
- Step 2. Obtain estimators $\hat{q}_1, \ldots, \hat{q}_m \in [0, 1]$, where $\hat{q}_j$ denotes an estimate of $P(H_j \text{ is null} \mid p_j, x_j), j = 1, \ldots, m$, taking values in $[\epsilon, 1]$ (if $\hat{q}_j < \epsilon$, let $\hat{q}_j = \epsilon$);
- Step 3. Apply the weighted BH procedure of Genovese *et al.* (2006) with weights $\hat{q}_j^{-1}$. Specifically, define
$$\hat{k} = \max\left\{k \geq 1 : p_j \leq \left(\frac{\alpha}{\hat{q}_j} \cdot \frac{k}{m}\right) \wedge \tau\right\}$$
  with the convention that $\hat{k} = 0$ if this set is empty.
- Step 4. Reject any $p_j$ satisfying $p_j \leq \left(\frac{\alpha}{\hat{q}_j} \cdot \frac{\hat{k}}{m}\right) \wedge \tau$ for a total of $\hat{k}$ rejections.

SABHA is a generalization of several methods including BH. Step 3 and 4 are equivalent to the BH procedure at significant level $\frac{\alpha}{\hat{q}_j}$. If we set $\tau = 1$ and $\hat{q}_j = 1$ for all $j = 1, \ldots, m$, then SABHA is exactly the original BH procedure.

#### A.4 OrderShapeEM

OrderShapeEM (Cao *et al.* , 2022) is a Lfdr based multiple testing procedure incorporating auxiliary information. A monotone relationship is imposed between the auxiliary covariates and the prior probabilities of being null to estimate the unknowns, which can be further used to estimate the Lfdr. Here we conduct a thought experiment to illustrate the monotone constraint. We are given data consisting of a pair of values $(p_j, x_j)_{j=1}^m$, where $p_j$ represents the primary $p$-value, $x_j$ represents corresponding auxiliary information, and they are independent conditional on the hidden true state $\theta_j$ for $j = 1, \ldots, m$. Suppose

$$p_j \mid \theta_j \overset{\text{ind}}{\sim} (1-\theta_j)U(p_j) + \theta_j f_1(p_j),$$
$$x_j \mid \theta_j \overset{\text{ind}}{\sim} (1-\theta_j)U(x_j) + \theta_j f_1(x_j),$$

where $\theta_j = 1$ if $H_j$ is alternative and $\theta_j = 0$ if $H_j$ is null, $U(\cdot)$ is the density function of $p$-values or auxiliary covariates under the null hypothesis and $f_1(\cdot)$ is the density function of $p$-values or auxiliary covariates under the alternative hypothesis. Suppose $P(\theta_j = 0) = \tau_0$ for all $j = 1, \ldots, m$. Using the Bayes rule and the conditional independence between $p_j$ and $x_j$

given $\theta_j, j = 1, \ldots, m$, we have the conditional distribution of $p_j \mid x_j$ as follows:

$$
\begin{aligned}
f(p_j \mid x_j) =& \frac{f(p_j, x_j \mid \theta_j = 0)\tau_0 + f(p_j, x_j \mid \theta_j = 1)(1 - \tau_0)}{f(x_j \mid \theta_j = 0)\tau_0 + f(x_j \mid \theta_j = 1)(1 - \tau_0)} \\
=& \frac{f(p_j \mid \theta_j = 0)f(x_j \mid \theta_j = 0)\tau_0 + f(p_j \mid \theta_j = 1)f(x_j \mid \theta_j = 1)(1 - \tau_0)}{f(x_j \mid \theta_j = 0)\tau_0 + f(x_j \mid \theta_j = 1)(1 - \tau_0)} \\
=& \frac{U(p_j)U(x_j)\tau_0 + f_1(p_j)f_1(x_j)(1 - \tau_0)}{U(x_j)\tau_0 + f_1(x_j)(1 - \tau_0)} \\
=& U(p_j)\gamma_0(x_j) + f_1(p_j)(1 - \gamma_0(x_j)),
\end{aligned}
$$

where

$$
\gamma_0(x) = \frac{U(x)\tau_0}{U(x)\tau_0 + f_1(x)(1 - \tau_0)} = \frac{\tau_0}{\tau_0 + \frac{f_1(x)}{U(x)}(1 - \tau_0)}.
$$

Under the monotone likelihood ratio assumption (Sun and Cai , 2007):

$$
f_1(p) \text{ is decreasing in } p,
$$

$\gamma_0(x)$ is a monotone increasing function of $x$. Therefore, the order of $x_j$ generates a ranked list of the primary hypotheses $H_1, \ldots, H_m$ through the conditional prior probability $\gamma_0(x)$. More details of the algorithm can be found in Cao *et al.* (2022)

### B Simulation details

#### B.1 Primary data generation

In each realistic simulation setting, based on the spatially resolved mouse olfactory bulb data and parameters inferred by SPARK, we simulated the primary gene expression data on $n = 260$ spatial spots. For each gene in turn, the read counts on spot $i (i = 1, \ldots, n)$ were simulated as follows:

$$y_i \sim \text{Poi}(N_i\lambda_i),\ \log\lambda_i = \beta + \epsilon_i, \tag{S1}$$

where $\beta$ is a scalar representing the intercept as the dimension of $\mathbf{x}_i$ is $k = 1$. We want to simulate $y_i$ from $y_i \sim \text{Poi}(N_i\lambda_i)$, where $N_i$ denotes the total read counts of all genes on spot $i$ and can be obtained from the mouse olfactory bulb data. The underlying relative gene expression level $\lambda_i$ is unknown and can be generated from (S1). The error term $\epsilon_i$ measures the random noise independent of spatial locations. So it follows the same distribution for SE and non-SE genes, namely $\epsilon_i \sim N(0, \tau_2)$. We specified the value of $\tau_2$ to be 0.35, which is the median of estimates for different genes inferred from SPARK. The intercept $\beta$, which represents the mean of $\log\lambda_i$, takes different values for SE and non-SE genes. If the focused gene is a non-SE gene, we set $\beta = -10.2$ for $i = 1, \ldots, n$, which is shared across all spots and corresponds to the median of the intercept estimated by SPARK in the mouse olfactory bulb data. On the other hand, if the gene to be simulated is a SE gene, the spatial differential expression pattern is introduced by dividing the $n$ spots into two groups based on spatial Pattern I illustrated in Sun *et al.* (2020): a group of spots with low expression levels and a group of spots with high expression level. Then for the spots in the low expression group, $\beta = -10.2$; and for the spots in the high expression group, we specified the fold change to be 3, i.e., $\beta$ was set to be three-fold higher than $-10.2$ on rate parameter scale (i.e., $e^{\beta} = 3 \cdot e^{-10.2}$). Finally, for each gene in turn, we generated $\lambda_i$ through (S1), and $y_i$ was generated from $\text{Poi}(N_i\lambda_i)$.

In each setting, we simulated $y_i, i = 1, \ldots, n$ for all $m$ genes one by one, and then used SPARK to analyze the synthetic spatial transcriptomic data to obtain primary $p$-values for SE identification.

#### B.2 Auxiliary data generation

We used the two-sample RNA-seq data to provide auxiliary information for the SE analysis. RNA-seq read counts for $m = 10,000$ genes from two groups each consisting of 50 samples were generated based on the Negative Binomial (NB) distribution. In each setting, for gene $i$ in group $j$ $(j = 1, 2)$, its read count was simulated from $\text{NB}(\mu_{ij}, \phi_{ij})$, where $\mu_{ij}$ is the mean parameter and $\phi_{ij}$ denotes the dispersion parameter. To perform more realistic RNA-seq simulations, we used the *generateSyntheticData* function from R package *compcodeR* v1.28.0 (Soneson , 2014) to generate the synthetic count data with mean and dispersion parameters estimated from RNA-seq datasets (Pickrell *et al.* , 2010; Cheung *et al.* , 2010). In accordance with the primary spatial data, $\psi$ ($\psi = 7.5\%, 10\%$ or $15\%$) of the $m$ genes were simulated to be differentially expressed between the two groups (equally distributed between up- and down-regulated in group 2 compared to group 1).

The mean and dispersion parameters for each gene in the two groups can be generated based on the specified up- and down-regulated genes. Specifically, $\mu_{i1}$ and $\phi_{i1}$ $(i = 1, \ldots, m)$ were obtained by pair-wise sampling from the parameters estimated from real data. $\phi_{i2} = \phi_{i1}$, $\mu_{i2}$ was simulated by the product of $\mu_{i1}$ and the fold change ($\text{FC}_i$). For non-differentially expressed genes, $\text{FC}_i = 1$. For differentially expressed genes, we pre-set a baseline effect size 1.5, then the effect size of each gene was obtained by adding a random variable from the exponential distribution (with rate 1). Then $\text{FC}_i$ for up- and down-regulated genes were set as the effect size and the inverse of the effect size, respectively. Finally, with the simulated $\mu_{ij}$ and $\phi_{ij}$ for $i = 1, \ldots, m$ and $j = 1, 2$, the counts for $m$ genes across samples in two groups can be simulated from $\text{NB}(\mu_{ij}, \phi_{ij})$. We filtered the dataset by excluding the genes with zero counts in all samples (i.e., those for which the total count is 0).

The informativeness was introduced by the consistency of the non-null structures between the auxiliary and primary datasets. For auxiliary sequence, when $\psi = 7.5\%$, there are 750 non-null genes in total, if we randomly select 750 genes from the first $1,000$ genes to be differentially expressed, it would produce more informative auxiliary covariates than we randomly select 750 non-null genes from the first $1,500$ genes. We explored the effect of the covariate sequence under three different informative levels (i.e., "uninformative", "weak" and "strong") on the performance of the primary hypothesis testing. The uninformative covariate sequence can be simply generated by randomly sampling from a standard uniform distribution $\mathcal{U}(0,1)$. To generate a weakly informative covariate sequence, the differentially expressed genes in the auxiliary dataset were randomly selected from all $m = 10,000$ genes. As to the strongly informative covariates, for corresponding SE gene proportion of $\psi = 7.5\%, 10\%$ and $15\%$, we randomly selected 750, $1,000$ and $1,500$ genes from the first $1,500$, $2,000$ and $3,000$ genes to be differentially expressed, respectively.

After obtaining the auxiliary RNA-seq dataset, the R package *DESeq2* v1.30.1 (Love *et al.* , 2014) was applied to perform differential analysis, and the resulted $p$-values were utilized as auxiliary covariates for statistical analysis of SE genes.

### C Details of the real data acquisition

We implemented the proposed integrative analysis method on four published spatial transcriptomic RNA-seq datasets incorporating auxiliary information obtained from multi-omics studies. The raw spatial transcriptomic count data of mouse olfactory bulb and human breast cancer (Ståhl *et al.* , 2016) used in the case studies were originally downloaded from `https://www.spatialresearch.org/resources-published-datasets/doi-10-1126science-aaf2403`; the raw mouse cerebellum dataset was available at the Broad Institute's single-cell repository (`https://singlecell.broadinstitute.org/single_cell/`) with ID SCP948; and the human heart data was collected from the 10X Visium spatial gene-expression repository. The website is `https://support.10xgenomics.com/spatial-gene-expression/datasets/`

`1.1.0/V1_Human_Heart`. For SPARK, we filtered out genes that were expressed in less than 10% of the spatial locations and selected spatial locations with at least 10 total read counts to obtain the final primary datasets.

#### C.1 Mouse olfactory bulb data

The first dataset was from the mouse olfactory bulb data, consisting of $11,274$ genes measured on 260 spots. We used the R package *SPARK* v1.1.2 to make SE analysis and obtained the primary $p$-values. The auxiliary scRNA-seq dataset is publicly available at `http://mousebrain.org/adolescent/tissues.html` (Zeisel *et al.* , 2018), consisting of $27,998$ genes measured on $20,636$ mouse olfactory cells. We used the R package *Seurat* v4.0.2 (Satija *et al.* , 2015; Hao *et al.* , 2021) to analyze the scRNA-seq data and tested for differentially expressed features, resulting in $43,724$ $p$-values for genes in each cluster compared to all remaining cell-types. By filtering out duplicate genes in multiple clusters and only keeping the smallest $p$-value across all involved clusters for each gene, we obtained an auxiliary $p$-value sequence consisting of $8,084$ genes.

Then we matched the primary and auxiliary $p$-values by gene, and took the $7,026$ pairs of $p$-values as input to the subsequent multiple testing procedures.

#### C.2 Mouse cerebellum data

After obtaining the processed primary mouse cerebellum dataset composed of $20,117$ genes measured on $11,626$ beads from Zhu *et al.* (2021), we further filtered out genes following the default setting of SPARK, resulting in a final set of dataset containing 523 genes on $11,626$ beads. Then we used the R package *SPARK* v1.1.2 to make SE analysis and obtained the primary $p$-values. The auxiliary scRNA-seq dataset of mouse cerebellum is available at `http://mousebrain.org/adolescent/tissues.html` (Zeisel *et al.* , 2018), which contains

27,998 genes measured on 12,312 mouse cerebellum cells. The R package *Seurat* (Satija *et al.* , 2015; Hao *et al.* , 2021) was applied to analyze the scRNA-seq data, yielding $p$-values of 23,317 genes for 15 cell-type clusters compared to all remaining cell-types. We filtered out duplicate genes found in multiple clusters, and only kept the smallest $p$-value for each gene. Finally, an auxiliary $p$-value sequence for 6,666 genes was obtained.

Then we matched the primary and auxiliary $p$-value sequences by gene and used the paired $p$-values of 502 genes common to both studies as input to OrderShapeEM.

#### C.3 Human breast cancer data

Table S1: Seven different types of auxiliary $p$-value sequences obtained by free combination of three auxiliary datasets. In each case, the symbol "+" and "−" indicate the inclusion and exclusion of corresponding auxiliary dataset, respectively.

| | TCGA BRCA | TCGA THCA | Gene-based GWAS |
|---|---|---|---|
| Case 1 | + | − | − |
| Case 2 | − | + | − |
| Case 3 | − | − | + |
| Case 4 | + | − | + |
| Case 5 | + | + | − |
| Case 6 | − | + | + |
| Case 7 | + | + | + |

The third dataset was from the study of human breast cancer, which contained read counts of 5,262 genes measured on 250 spatial spots. We used the R package *SPARK* v1.1.2 to make SE analysis of this dataset and obtained the primary $p$-values of 5,262 genes. For each gene, we created the auxiliary covariates from three different studies: The Cancer Genome Atlas (TCGA) RNA-seq data from the Breast Cancer program (BRCA) consisting of 55,112 genes, TCGA RNA-seq data from the Thyroid Cancer program (THCA) consisting of 39,788 genes, and GWAS summary data from the Breast Cancer Association Consortium (BCAC) for women of European ancestry consisting of $11,089,342$ SNPs (`http://bcac.ccge.medschl.cam.ac.uk/bcacdata/oncoarray`) (Michailidou *et al.* , 2017). We utilized TCGA THCA RNA-seq data as auxiliary information by virtual of the pleiotropy between breast and thyroid cancers. The two TCGA RNA-seq datasets were analyzed with R package *DESeq2* v1.30.1 (Love *et al.* , 2014). For the GWAS summary statistics, we conducted a gene-based association analysis using the aggregated Cauchy association test (ACAT) with the R package *sumFREGAT* v1.2.1 (Svishcheva *et al.* , 2019; Liu *et al.* , 2019) to obtain auxiliary $p$-values of $18,871$ genes.

The three auxiliary datasets have seven different types of auxiliary $p$-value sequences through free combinations (Table S1). There were $4,801$ overlapping genes in the primary study and the three auxiliary studies. In each case, we took the primary and corresponding auxiliary $p$-values of these $4,801$ genes as input to make integrative analysis.

#### C.4 Human heart data

10X Genomics commercialize and improve the original spatial transcriptomics in their Visium Spatial Gene Expression platform with higher resolution (55 $\mu$m per spot). The Visium spatial gene expression dataset of human heart contains $36,601$ genes measured on $4,247$ spots. We filtered out mitochondrial genes and genes that were not expressed in any spot and obtained a final set of $20,904$ genes on $4,247$ spots. For the auxiliary information, we downloaded a publicly available combined scRNA-seq and single nuclei RNA-seq (snRNA-seq) dataset of 485K cardiac cells (Litviňuková *et al.*, 2020) at `https://cells.ucsc.edu/?ds=heart-cell-atlas` and performed differential expression analysis between 27 different cell-type clusters with R package *Seurat* v4.0.2 (Satija *et al.* , 2015; Hao *et al.* , 2021), yielding $43,673$ $p$-values. We only kept the smallest $p$-value of the same gene among all recurring $p$-values, and finally obtained an auxiliary $p$-value sequence of $4,088$ genes.

By taking the intersection of genes in the primary study and auxiliary study, we obtained $3,848$ genes in both studies. Then we performed SPARK and SPARK-X (Zhu *et al.* , 2021) on the $3,848$ genes measured on $4,247$ spots in the Visium human heart dataset to produce primary $p$-values. Finally, we took $3,848$ pairs of $p$-values as input to OrderShapeEM.

### D Supplementary results

In this section, we present supplementary analysis and validation results of the four case studies to provide more biological evidence and interpretations for the discoveries of our proposed integrative analysis framework.

#### D.1 Mouse olfactory bulb data

We used the UMAP method implemented in R package *umap* v0.2.8.0 to reduce the $1,430$ significant SE genes identified by OrderShapeEM into two dimensions. With the cell labels obtained from hierarchical agglomerative clustering using R package *amap* v0.8-18, the cell distribution of those genes were depicted in Fig. S1(A), most genes can be grouped into three cell-type clusters. OrderShapeEM additionally identified 619 SE genes compared to SPARK and 317 SE genes compared to BH. We clustered these uniquely identified SE genes into three groups and obtained three major spatial expression patterns corresponding to the mitral cell layer, the glomerular layer and the granular layer in mouse olfactory bulb. (Fig. S1(B)(C)). We randomly selected six SE genes identified by OrderShapeEM but not by BH and plotted their spatial expression patterns in Fig. S1(D). We also calculated the Moran's $I$ statistics (Moran, 1950) for SE genes detected by OrderShapeEM and the original SPARK as well as for all genes in the dataset to provide an additional validation of the spatial auto-correlation. Fig. S1(E) shows the boxplot result, Moran's $I$ statistic of SE genes detected by OrderShapeEM is generally larger than the Moran's $I$ statistic of all genes in the dataset. SPARK has larger

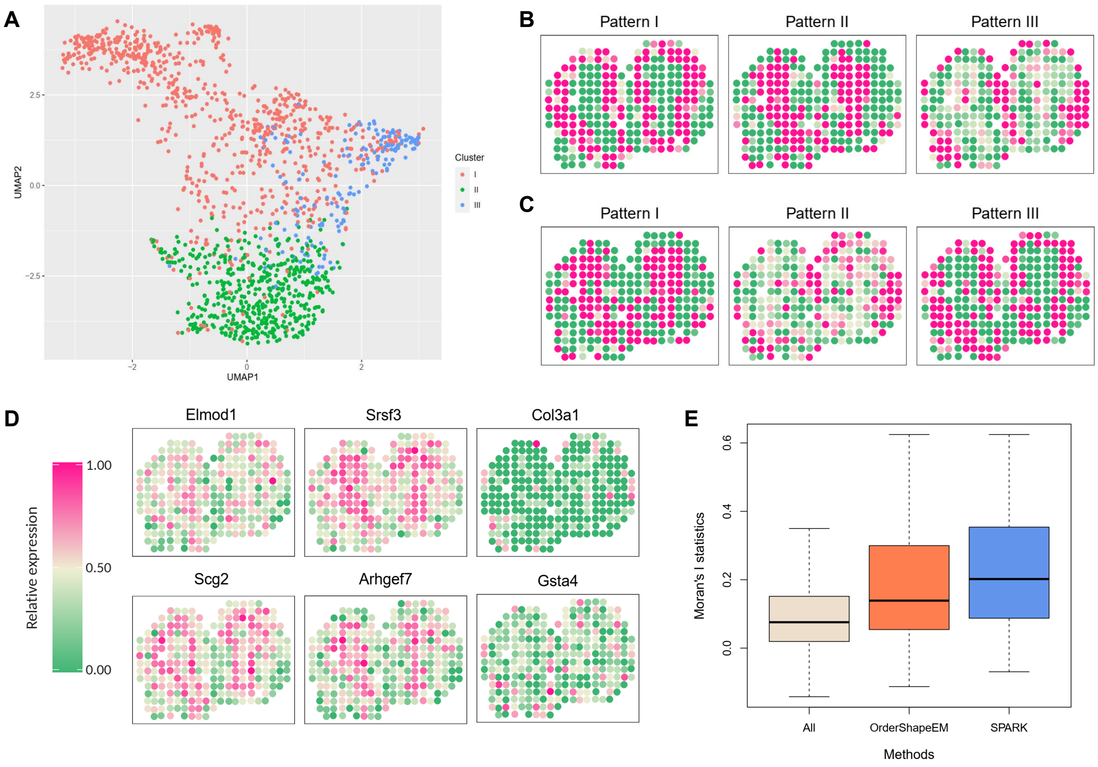


Figure S1: Analysis and validation results of the mouse olfactory bulb data. (A) Scatter plot visualizes the spatial clusters of the SE genes identified by OrderShapeEM. (B) Spatial expression patterns summary based on the 691 SE genes identified by OrderShapeEM but not by SPARK when the number of clustering is set to be three. (C) Spatial expression patterns summary based on the 317 SE genes identified by OrderShapeEM but not by BH. (D) Spatial expression patterns for six SE genes identified by OrderShapeEM but not by BH. (E) Boxplot of Moran's $I$ statistics for different methods and all genes in the dataset.

Moran's $I$ statistic than OrderShapeEM as it is conservative.

#### D.2 Mouse cerebellum data

We clustered the 459 significant SE genes identified by OrderShapeEM in the mouse cerebellum data into two groups using the hierarchical agglomerative clustering algorithm. With the cell labels obtained, we reduced the dimension of the SE genes using the UMAP method and visualized the clusters in Fig. S2(A). The comparison of the Moran's $I$ statistics for the 144 SE genes additionally detected by OrderShapeEM compared to genes not identified by

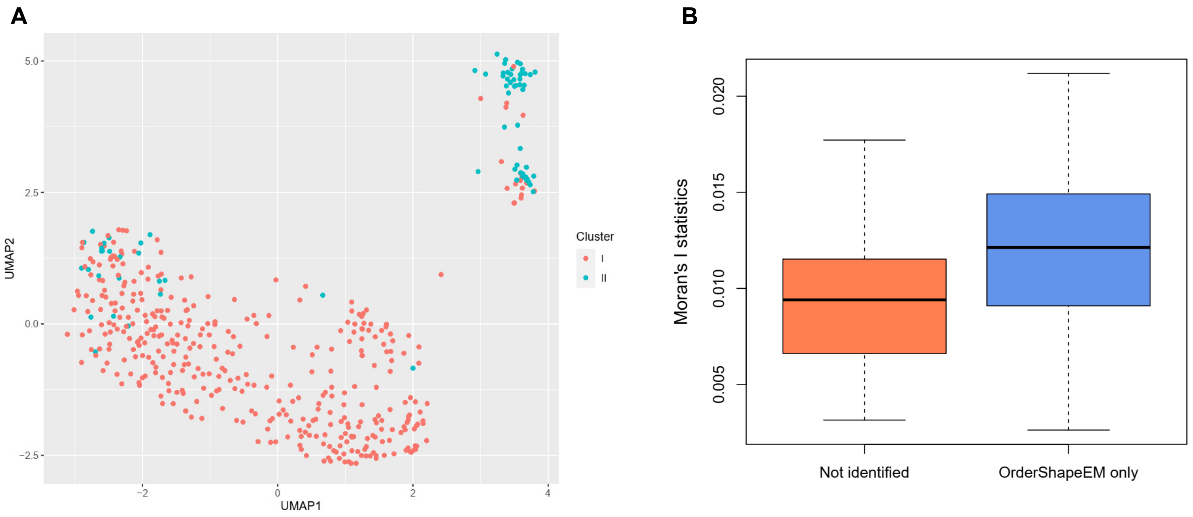


Figure S2: Analysis and validation results of the mouse cerebellum data. (A) Clustered scatter plot of SE genes identified by OrderShapeEM. (B) Boxplot of Moran's $I$ statistics for SE genes uniquely identified by OrderShapeEM and genes not identified by OrderShapeEM.

OrderShapeEM is presented in Fig. S2(B). We see higher overall spatial auto-correlation in genes additionally identified by OrderShapeEM.

#### D.3 Human breast cancer data

In all seven cases using different combinations of auxiliary information (Table S1), we clustered the significant SE genes identified by OrderShapeEM on the FDR cutoff of 0.05 into two groups. As in Fig. S3(B), most SE genes detected by OrderShapeEM can be grouped into two clusters corresponding to the spatial expression patterns. After excluding the SE genes identified by SPARK, the additional SE genes detected by OrderShapeEM can still be summarized to two distinct spatial expression patterns. Fig. S3(A) shows the summarized spatial patterns in Case 7 which incorporated the combination of three auxiliary studies, and Fig. S4 plots spatial expression patterns of 20 genes randomly selected from the SE genes only identified by OrderShapeEM. Furthermore, we calculated Moran's $I$ statistics for SE genes identified by OrderShapeEM and SPARK as well as all genes in the dataset. As shown in

Fig. S3(C), in all cases, Moran's $I$ statistics for the SE genes identified by OrderShapeEM is higher than Moran's $I$ statistics of all genes tested.

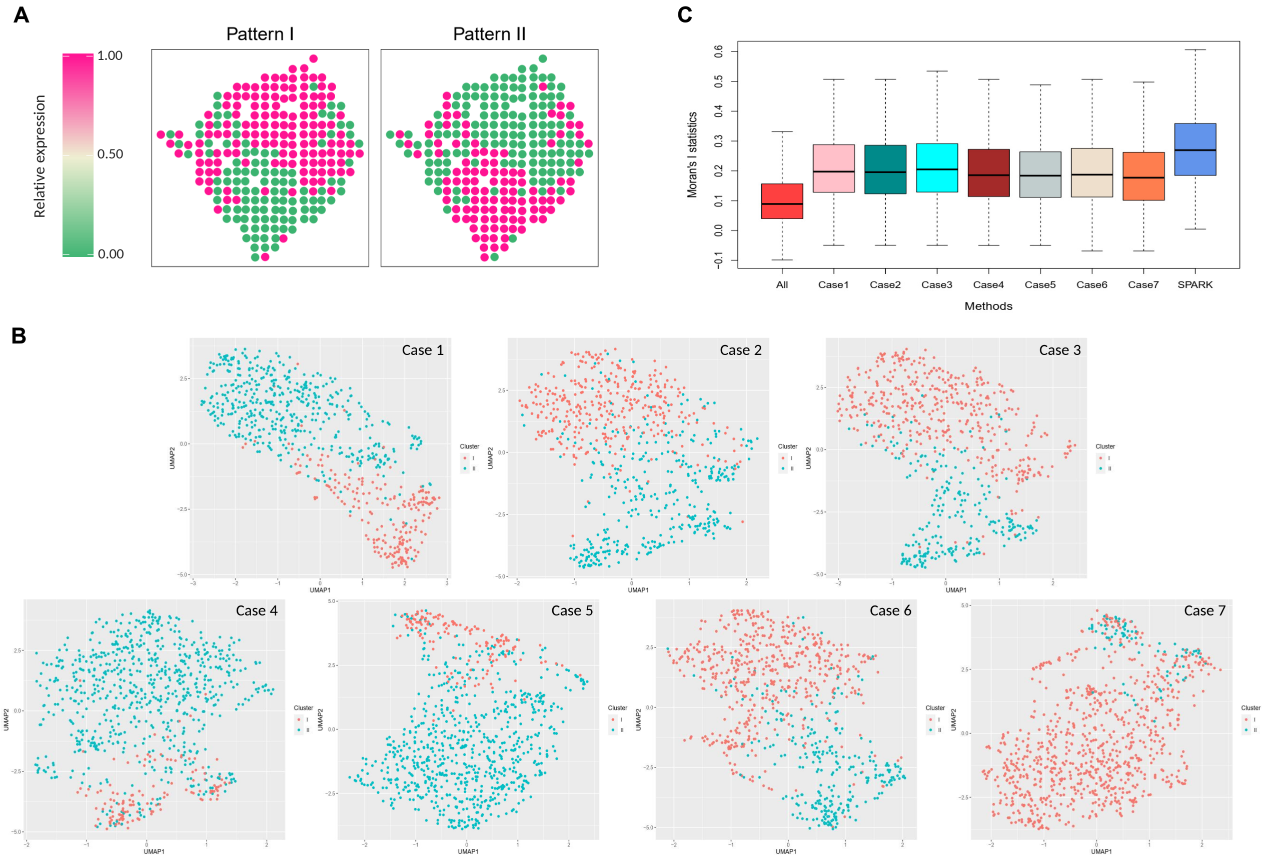


Figure S3: Analysis and validation results of the human breast cancer data. (A) Spatial expression patterns summarized based on the 638 SE genes identified by OrderShapeEM (Case 7) but not by SPARK when the number of clustering is set to be two. (B) Clustered scatter plot of SE genes identified by OrderShapeEM in all seven cases. (D) Boxplot of Moran's $I$ statistics for different methods and all genes in the dataset. Case1 - Case7 represent OrderShapeEM incorporating different auxiliary covariates.

#### D.4 Human heart data

SPARK performs well for spatial transcriptomic data measured on a moderate number of spatial locations (e.g., hundreds or thoudands). The computational complexity of SPARK scales cubically with respect to the number of spots. Consequently, it takes hours to days for

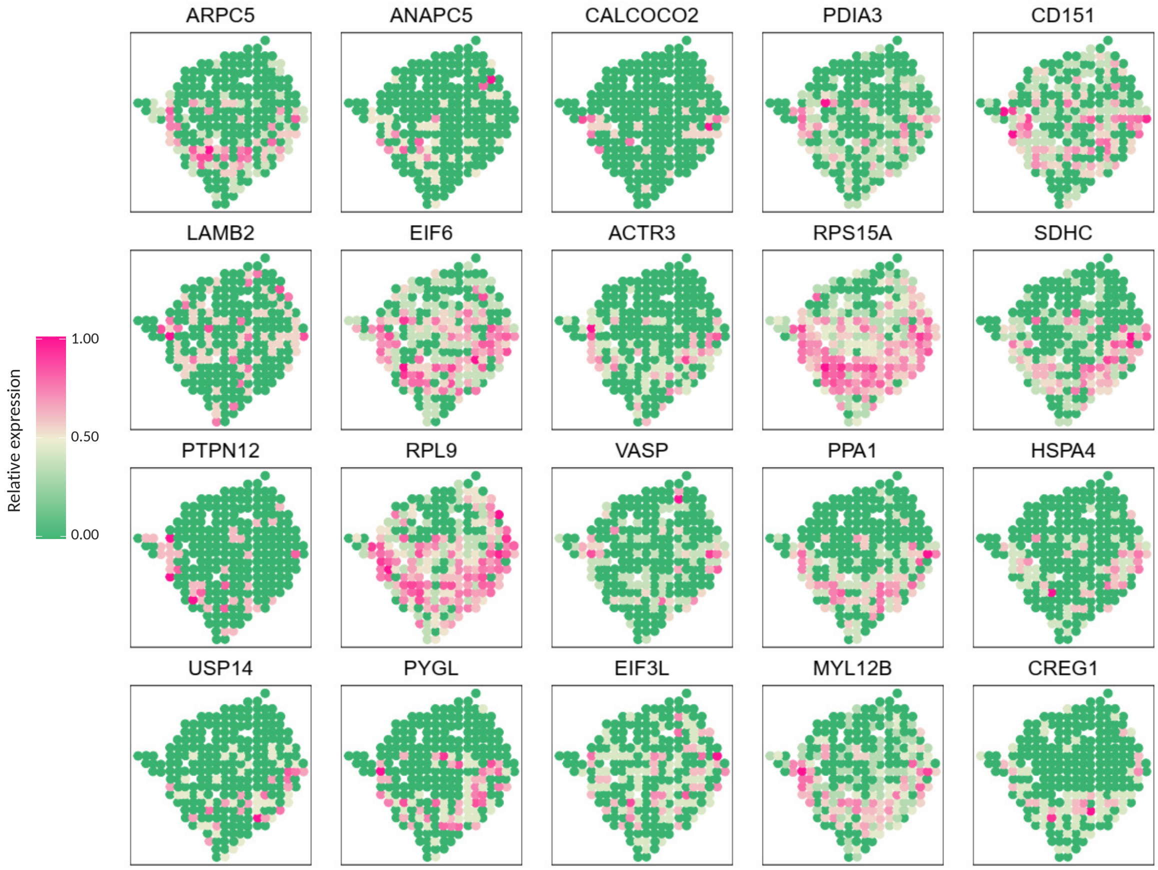


Figure S4: Spatial expression patterns of 20 genes randomly selected from the SE genes identified by OrderShapeEM (Case 7) but not by SPARK in the human breast cancer data.

SPARK to analyze large-scale spatial transcriptomic data (e.g., with $\geq 3,000$ spatial spots) (Sun *et al.* , 2020). We also implemented a scalable non-parametric method, SPARK-X (Zhu *et al.* , 2021), in the proposed integrative analysis framework to analyze the large-scale Visium human heart data. Both SPARK and SPARK-X can produce calibrated $p$-values (Sun *et al.* , 2020; Zhu *et al.* , 2021). As in Fig. S5(A), the quantile-quantile plots of $p$-values for the human heart data suggests different power of SPARK and SPARK-X.

Here we summarize the detailed analysis results of the human heart data by implementing SPARK-X. Using the $p$-values of $3,848$ genes produced by SPARK-X as primary $p$-values, the original SPARK-X (BY) identified 488 SE genes. By incorporating corresponding auxiliary $p$-values resulted from the combined scRNA-seq and snRNA-seq data in human heart tissues,

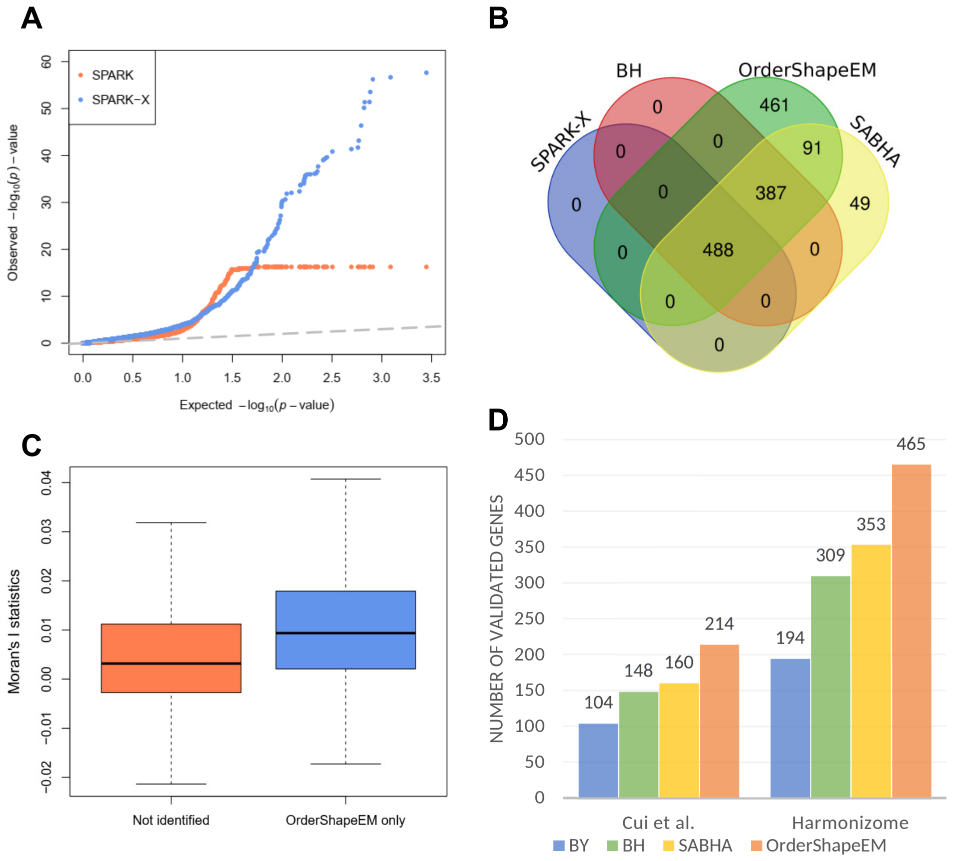


Figure S5: Analysis and validation results of the human heart data by implementing SPARK-X. (A) Quantile-quantile plot of the observed $-\log_{10}$ $p$-values from the SPARK and SPARK-X methods against the expected $-\log_{10}$ $p$-values under the null in the human heart data. (B) Venn diagram shows the number of SE genes identified by different methods based on SPARK-X at a target FDR level $\alpha = 0.05$ and their resulting discovery set intersections. (C) Boxplot displays Moran's $I$ statistics for the SE genes uniquely identified by OrderShapeEM and genes not detected by OrderShapeEM. (D) Barplot displays the number of SE genes identified by each method that were validated in two reference gene lists: one from (Cui *et al.*, 2019) and the other from the Harmonizome database.

OrderShapeEM identified $1,427$ significant SE genes on an FDR cutoff of 0.05, including all SE identified by SPARK-X. The Venn diagram in Fig. S5(B) shows the number of discoveries and set intersections of different methods at an FDR cutoff of 0.05. We calculated the Moran's $I$ statistics to evaluate the spatial auto-correlation of the SE genes identified by OrderShapeEM but not by SPARK-X. The boxplot result is presented in Fig. S5(C). We also provided two additional lines of evidence to validate the results of the framework based on SPARK-X using two set of genes from previous literature (Cui *et al.*, 2019) and Harmonizome database (Rouillard *et al.*, 2016). As shown in Fig. S5(D), 110 and 271 SE genes additionally identified by OrderShapeEM are successfully validated in the two reference gene lists, respectively.

Table S2 summarizes the computation time in seconds of different methods for analyzing four datasets. Computations were carried out in an Intel(R) Core(TM) i7-9750H 2.6GHz CPU with 64.0 GB RAM laptop with 10 threads for SPARK and 1 thread for SPARK-X.

Table S2: Computation time (in seconds) of different methods for analyzing the four datasets: the mouse olfactory bulb (MOB), the mouse cerebellum (MC), the human breast cancer (HBC) and the human heart (HH).

| Dataset | # of Genes /Samples | SPARK | SPARK-X + OrderShapeEM | SPARK-X | SPARK-X + OrderShapeEM |
|---|---|---|---|---|---|
| MOB | 7026/260 | 410.28 | 412.34 | 14.12 | 15.75 |
| MC | 502/11625 | 53119.01 | 53119.20 | 1.12 | 1.20 |
| HBC | 4801/250 | 320.23 | 321.84 | 5.45 | 5.81 |
| HH | 3848/4247 | 11252.86 | 11253.75 | 11.87 | 12.31 |
